# Bridging the microstructural gap in human connectomics using hierarchical phase-contrast tomography as a reference for diffusion MRI in the human brain

**DOI:** 10.64898/2026.04.02.715729

**Authors:** Eric Wanjau, Matthieu Chourrout, Chiara Maffei, Yael Balbastre, Andrew Keenlyside, Joseph Brunet, Aikta Sharma, Susie Y. Huang, Paul Tafforeau, Bruce Fischl, Anastasia Yendiki, Peter D. Lee, Claire L. Walsh

**Author notes:** Contributing authors.

## Abstract

Diffusion MRI (dMRI) allows us to image the human connectome non-invasively, yet it provides indirect estimates of axonal orientations based on the diffusion of water molecules in millimeter-scale voxels, hence struggling to resolve complex micrometer-scale fiber geometries. Invasive methods for imaging axonal orientations ex vivo, e.g. histology, are destructive and limited to small volumes, creating a critical need for a non-destructive modality for imaging microscopic fiber orientations in 3D. Here, we use Hierarchical Phase-Contrast Tomography (HiP-CT) to characterize white matter architecture at the microscale. Applying structure-tensor analysis to HiP-CT data, we compute fiber Orientation Distribution Functions and perform tractography analogous to dMRI. Across multiple brain regions, HiP-CT derived fiber architecture shows strong correspondence with that derived from dMRI while revealing substantially greater microstructural complexity. Despite its label-free nature, we demonstrate that vascular structures minimally confound HiP-CT orientation estimates. These results establish HiP-CT as a reference microscopic modality that can complement dMRI in multi-scale studies of white-matter organization.

## 1 Introduction

The function of the nervous system is intricately shaped by the structure, arrangement and interconnection of neurons and neuronal populations [1, 2]. Capturing this organization across spatial scales, ranging from cellular microstructure to long-range fiber architecture, remains fundamental to understanding brain function, development, and disease processes.

Hierarchical Phase-Contrast Tomography (HiP-CT) has recently emerged as a powerful imaging modality capable of addressing this challenge[3]. HiP-CT exploits X-ray phase contrast, in which local variations in electron density rather than absorption, provide contrast. Leveraging the Extremely Brilliant Source of the European Synchrotron Radiation Facility (ESRF-EBS), HiP-CT delivers isotropic, three-dimensional imaging of intact human brains. Whole-brain volumetric overviews are followed by targeted re-imaging of selected brain regions at progressively higher spatial resolutions. Crucially, these high-resolution volumes are intrinsically spatially aligned to the whole-brain dataset, as all acquisitions are performed within the same intact specimen and common coordinate frame. X-ray phase contrast provides sensitivity to subtle density differences in soft biological tissues, enabling visualization of white-matter fiber organization, as well as vasculature, and subcortical nuclei [4]. HiP-CT has been used to image intact human brain spanning whole-organ overviews at 20 −7 µm/voxel down to individual specialized cells, including Purkinje cells in the cerebellum at 6.5 −1.3 µm/voxel [3–5].

Within the human brain, white matter represents a particularly compelling target for HiP-CT. White matter is composed of densely packed, coherently oriented axonal bundles surrounded by other tissue components such as glial cells and vasculature[1, 2]. HiP-CT’s phase-contrast and multi-scale capability positions it as a critical tool for resolving this complex microenvironment, and presents a unique opportunity to bridge the gap between macroscale connectomes obtained from in vivo neuroimaging and nanoscale circuits reconstructed with electron microscopy.

Diffusion-weighted Magnetic Resonance Imaging (dMRI) is the method of choice for imaging white matter architecture in vivo and non-invasively. It infers axonal orientations by sensitizing the MR signal to the diffusion of water molecules in different directions [6–8]. Over the past decade, the Human Connectome Project (HCP) ushered in an era of significant innovations in dMRI acquisition techniques, with the introduction of MRI scanners with ultra-high gradient strength [9–13]. These developments have led to substantial improvements in dMRI resolution and signal-to-noise-ratio, allowing the estimation of fiber orientations with greater fidelity than ever before. To this day, dMRI has been used widely to studying white-matter organization and its disruption in neurological and psychiatric disorders [14, 15], including Alzheimer’s disease [16], Parkinson’s disease [17], and schizophrenia [18, 19].

While dMRI is undeniably a powerful clinical and research tool, its accuracy is limited by the indirect nature of its measurements, the modeling assumptions used to estimate orientations from these measurements, as well as its limited spatial resolution [6, 20]. Even the highest resolutions feasible with dMRI in the human brain are on the order of 100s of micrometers, whereas axons have diameters in the micron range (0.16 − 9 µm) [21]. As a result, there is growing interest in the ex vivo validation of dMRI against microscopy. Optical imaging, histology, and related synchrotron-based X-ray techniques have increasingly been adopted for this purpose, driven by advances in tissue preparation and labeling, automated imaging platforms, coherent radiation sources, and increased computational power [20, 22]. Validation of dMRI-derived fiber orientations has been performed using label-free optical methods such as three-dimensional polarized light imaging [23, 24], polarization-sensitive optical coherence tomography [7, 25, 26], and lightsheet microscopy in cleared tissue [20, 27]. While optical techniques provide high-resolution images (∼ 1 − 200 µm) capable of resolving white-matter fibers, they are limited by shallow imaging depth, reliance on physical sectioning, and the computational burden associated with reconstructing large volumetric datasets. Tissue-clearing approaches are further constrained by the penetration of clearing agents and stains in large samples [28]. As a result, scalability to whole-brain imaging remains a critical unmet challenge.

Synchrotron X-ray tomography addresses many of these limitations by enabling non-destructive, isotropic imaging of intact human organs at micron-level resolution [3, 22, 29–32]. As X-rays can easily penetrate through centimeters of soft tissue there is no need for physical sectioning to image deep structures. Synchrotron-based imaging thus holds the promise of probing white matter architecture down to the microscale across the entire brain, and thus serving as a reference modality for dMRI. HiP-CT is ideal for this purpose among synchrotron-based techniques, thanks to its compatibility with MRI scanning, its ability to image whole, intact organs without physical sectioning and its hierarchical, multi-scale imaging strategy. In this work, we demonstrate the utility of HiP-CT as a non-destructive, isotropic, three-dimensional modality for imaging white-matter architecture that is complementary to dMRI. Specifically, we show that:

1. HiP-CT can directly visualize white-matter architecture at a level of anatomical detail unattainable with dMRI.
2. Structure tensor analysis applied to HiP-CT data enables extraction of white-matter fiber orientation estimates and tractography reconstructions analogous to dMRI.
3. Vascular structures, which are clearly visible in HiP-CT data, exert minimal confounding effects on HiP-CT–based white-matter analysis.

Together, these findings position HiP-CT and dMRI as complementary modalities for multi-scale investigation of human brain structure. By integrating high-resolution X-ray phase-contrast imaging with dMRI, this work paves the way for multi-modal, multi-scale studies of brain connectivity in health and disease.

## 2 Results

### 2.1 HiP-CT can directly visualize brain microstructure

Figure 1 demonstrates the contrast and sensitivity of HiP-CT to white matter microstructure. The internal capsule, the red nucleus, and the pons are highlighted as example areas with different tissue microenvironments containing highly myelinated bundles and/or gray matter. We juxtapose HiP-CT at 15 µm isotropic resolution with a normalized spherical average image from dMRI (average of volumes acquired with *b* = 10000 *s/mm*^2^, normalized by the average of volumes acquired with *b* = 0) at 800 15 µm isotropic resolution. The normalized spherical average dMRI volume exhibits contrast between white matter, where signal decays slower due to restricted diffusion in myelinated fiber bundles, and gray mattter, where signal decays faster due to less restriction.

**Fig. 1.**
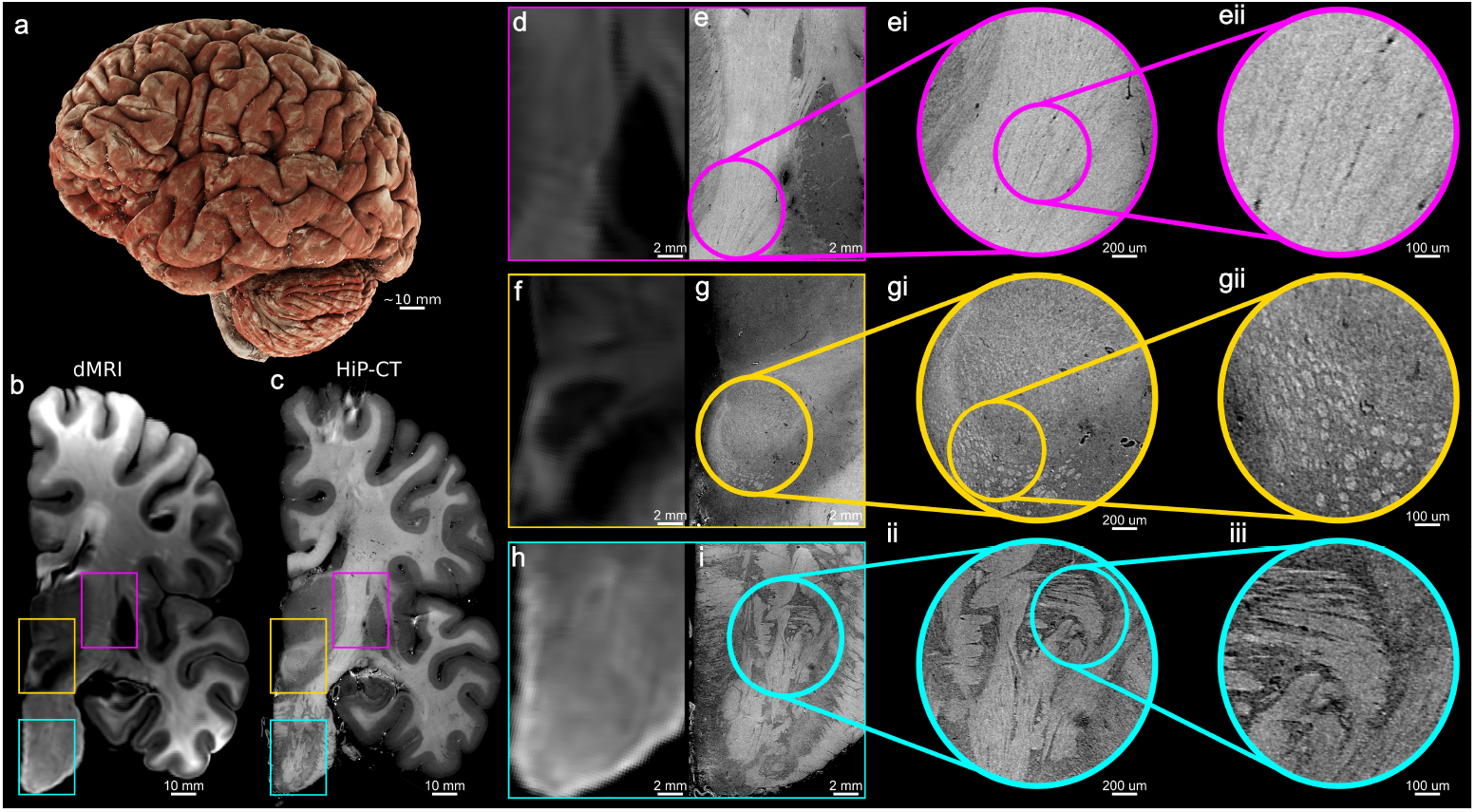
HiP-CT resolves fine white-matter microstructure. **a**, 3D volumetric rendering of the HiP-CT scanned hemisphere. **b** Coronal slice of the average diffusion-weighted image acquired at *b* = 10000 *s/mm*^2^, normalized by the average *b* = 0 image (800 µm isotropic) showing slower signal decay in white matter due to restriction. **c** Co-registered coronal HiP-CT slice (15 µm isotropic). **d-e Internal Capsule (Magenta):** In dMRI (d), the normalized spherical average signal can differentiate between white and gray matter but does not provide further contrast to resolve microstructure in white matter. HiP-CT (e) reveals the underlying microstructure as a tightly packed block of fascicles and demonstrates sharp anatomical boundaries between white matter and adjacent gray matter nuclei. The magnified insets (**ei, eii**) visualize the individual vertical striations of internal capsule. **(f, g) Red Nucleus (Yellow):** In dMRI (f), the normalized spherical average signal is low in the red nucleus due to isotropic diffusion within gray matter, while a surrounding bright rim indicates the presence of adjacent white matter fiber bundles. HiP-CT (g) resolves the “swirling” internal texture of the red nucleus. High-magnification insets (**gi, gii**) resolve fascicles ventral to the nucleus as discrete, bright, circular structures. **(h, i) Pons (Cyan):** dMRI (h) captures coarse anisotropic contrast of the transverse and longitudinal pontine tracts. HiP-CT (i) resolves the fine-scale interdigitation of these fiber systems, revealing discrete fascicles, local variations in orientation, and microvasculature. Zooms (**ii, iii**) resolve striations of transverse and vertical fiber bundles. Overall, HiP-CT provides direct imaging of microstructure that in dMRI can only be probed with further modeling of the signal.

#### The Internal Capsule

The internal capsule is a compact sheet of white matter that carries major ascending and descending fibers between the cerebral cortex and subcortical structures, including thalamic, subthalamic, and brainstem nuclei [33, 34]. The posterior limb of the internal capsule (**Figure 1 d**) has high signal intensity in the spherical average dMRI volume, as a region of highly restricted diffusion, consistent with a densely packed and coherent fiber pathway. HiP-CT (**Figure 1 e**) clearly reveals the striated boundary where white matter fibers enter into the adjacent gray matter regions of the thalamus and lentiform nucleus. The zoomed-in insets (**Figure 1 ei, eii**) further reveal that the internal capsule is not a homogeneous medium, but a tightly packed arrangement of parallel, vertically oriented fascicles.

#### The Red Nucleus

The red nucleus is a region characterized by a complex integration of gray and white matter, with afferent and efferent fiber bundles entering, exiting and coursing around it [35]. In the dMRI spherical average image (**Figure 1 f**), the red nucleus appears as a region of low signal intensity due to isotropic water diffusion within the gray matter. The rim of higher signal surrounding the red nucleus indicates the surrounding fiber bundles. The corresponding HiP-CT inset (**Figure 1 g**) resolves the internal structure of the red nucleus as a textured region consistent with the intra-nuclear confluence of fiber bundles. Ventral to the red nucleus, the zoomed HiP-CT insets (**Figure 1 gi, gii**) reveal a cluster of bright circular structures that correspond to bundles of axons. The distinct axonal bundles seen in HiP-CT are not directly visible in the dMRI signal, where further modeling is required to extract fiber orientations. In addition, HiP-CT concurrently resolves vessels that can be identified as tubular structures of low intensity.

#### The Pons

Anatomically, the pons is characterized by the interdigitation of descending longitudinal fibers and transverse fibers [36, 37]. In the pontine insets, the spherical average image from dMRI at 800 µm (**Figure 1 i**) shows high signal intensity, consistent with restricted diffusion, in structures corresponding to both the transverse and longitudinal fibers. HiP-CT (**Figure 1 h**) resolves the highly organized pattern of transverse pontine fibers interwoven with perpendicularly oriented fibers. Furthermore, HiP-CT directly resolves fine-scale local variations in fiber orientations, which in dMRI would require modeling to interrogate, as well as the presence of microvasculature (visible as dark tubular voids). In the high-resolution insets (**Figure 1 ii, iii**), fine-scale stratification of fiber bundles can be observed.

This comparison demonstrates that HiP-CT resolves white matter microstructure that dMRI can only infer indirectly, thus serving as a complementary reference modality for interpreting the results of downstream modeling on the dMRI signal.

### 2.2 Reconstructing white matter fiber architecture from HiP-CT with structure tensor analysis

Figure 2 illustrates the application of Structure Tensor Analysis (STA) to reconstruct fiber architecture from the 15 µm HiP-CT whole brain overview volume. This framework allows for the transformation of local intensity gradients into local distributions of fiber orientation. **Figure 2 a** shows a coronal slab of 52 slices which approximate a dMRI voxel in the *z* direction. HiP-CT resolves fiber bundles and their orientation as particularly evidenced in the pons inset **Figure 2 e** where a highly organized interweaving pattern of horizontally and vertically oriented fiber bundles is clearly delineated.

The primary output of STA is a voxel-wise map of eigenvectors that represent the principal fiber orientation. Using a standard Direction-Encoded Color map (Red: leftright, Green: anterior-posterior, Blue: superior-inferior), the resulting orientation map **Figure 2 b** reveals coherent, smoothly varying principal directions that are consistent with the observed HiP-CT data. The inset **Figure 2 f** demonstrates that the intensity gradient-based principal eigenvectors capture the dominant fiber orientation and reproduce the macro-anatomy of fiber bundles in the pons.

**Fig. 2.**
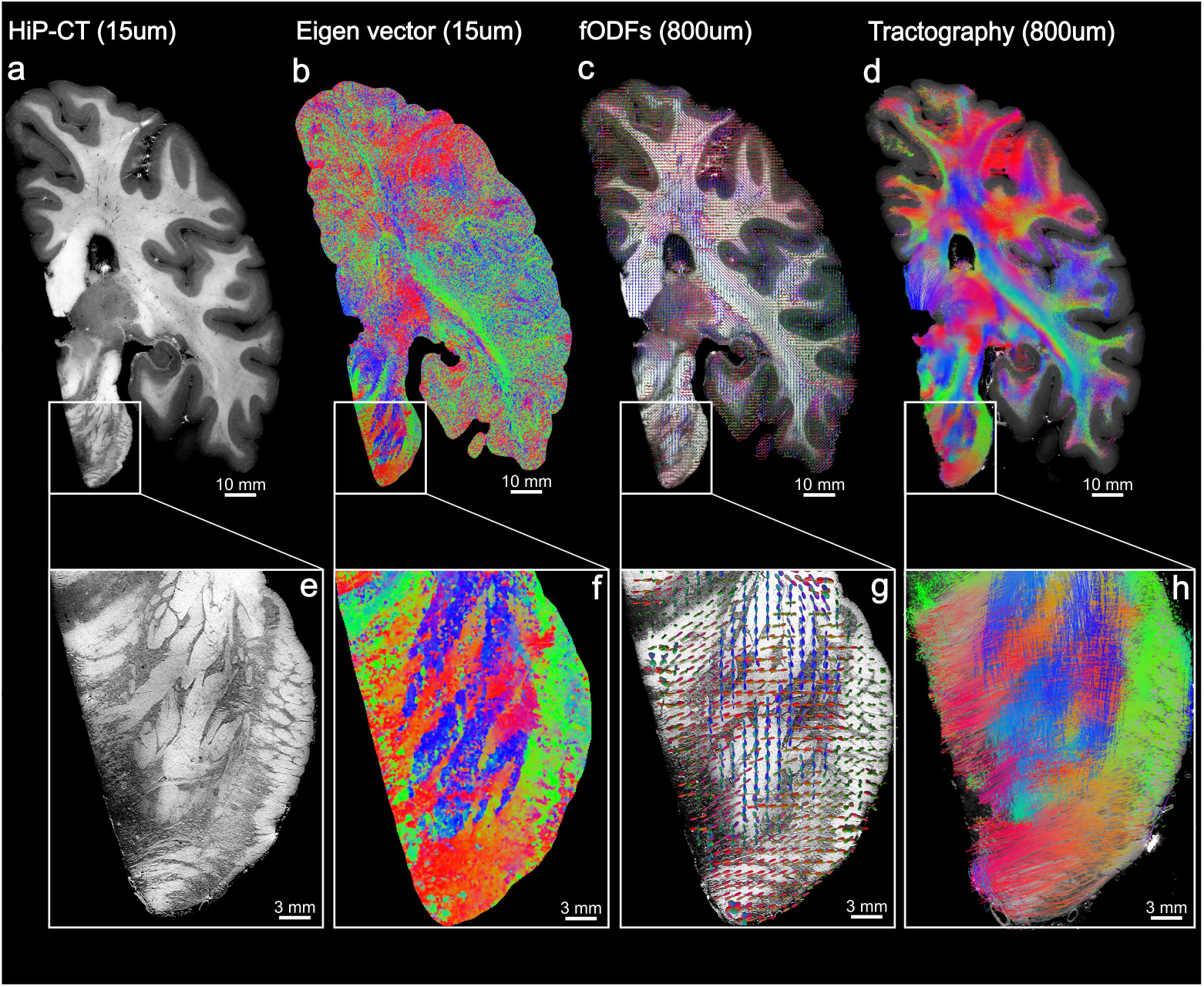
White matter fiber analysis workflow using Structure Tensor Analysis (STA) on HiPCT data. Starting with the tomographic HiP-CT intensity data at 15 µm (a), the pipeline computes a voxel-wise principal orientation map (b), where color indicates the dominant intensity gradient direction (Red: Left-Right, Green: Anterior-Posterior, Blue: Superior-Inferior). These local tensors are aggregated spatially into supervoxels of the same size as the dMRI voxel (800 µm) to estimate mesoscopic fiber Orientation Distribution Functions (fODFs) (c) visualized here as glyphs that model local fiber populations. Using the fODFs as a propagator, fiber trajectories can be reconstructed with probabilistic tractography. (d). The magnified insets of the pons (e–h) illustrate how this workflow resolves complex fiber geometries: the raw structural texture (e) yields an interdigitated orientation map (f) and multi-peaked fODF glyphs (g), which successfully represent the intricate crossing of vertical corticospinal and transverse pontocerebellar tracts (h).

To facilitate comparison with dMRI, the principal eigenvectors within dMRI-equivalent supervoxels (800 µm) were aggregated into spherical histograms, and fit with 8th order spherical harmonic functions. The resulting fiber Orientation Distribution Functions (fODFs) for the slab are visualized as 3D glyphs color-coded in a similar fashion as the principal orientation **Figure 2 c**. Zooming into the pons **Figure 2 g**, the fODF glyphs resolve the characteristic weaving of fiber bundles as seen in the raw HiP-CT images. The glyphs exhibit multiple distinct peaks, simultaneously capturing the descending longitudinal corticospinal tracts (blue vertical peaks) and the intersecting transverse pontocerebellar fibers (red horizontal peaks).

Using the fODFs as a propagator we performed probabilistic tractography, producing dense, continuous, anatomically plausible streamlines **Figure 2 d**. In the pontine inset, **Figure 2 h** the generated streamlines reconstruct a complex interweaving geometry, tracing the transverse fibers as they thread through the vertical fascicles.

Collectively, these results show that structure tensor analysis provides a robust framework for extracting continuous, anatomically meaningful white matter orientation information from HiP-CT. By accumulating these local tensors into voxel-wise fODFs, we establish a framework analogous to dMRI reconstruction methods, enabling direct, scale-matched comparative analysis to dMRI.

### 2.3 Qualitative comparison between dMRI and HiP-CT tractography at 800 µm

Comparison between Constrained Spherical Deconvolution (CSD)-based dMRI tractography (**Figure 3 a**) and STA-based HiP-CT tractography and (**Figure 3 b**), both at a resolution of 800 µm, shows broad global agreement in bundle organization. Coronal cross-sections (**Figure 3 c, d**) allows for direct evaluation of the finer structural details. HiP-CT tractography replicates the overall organization of major white matter structures previously established by dMRI and classical anatomical studies. Prominent fascicles such as the corticospinal tract (blue), the corpus callosum (red), and the cingulum bundle (green) occupy the same spatial footprints, and exhibit identical bulk orientations in both modalities (Figure 3 c, d). This global agreement provides evidence that local intensity gradients captured by HiP-CT structure tensor analysis provide a reliable means for reconstructing white matter fiber bundle orientation.

**Fig. 3.**
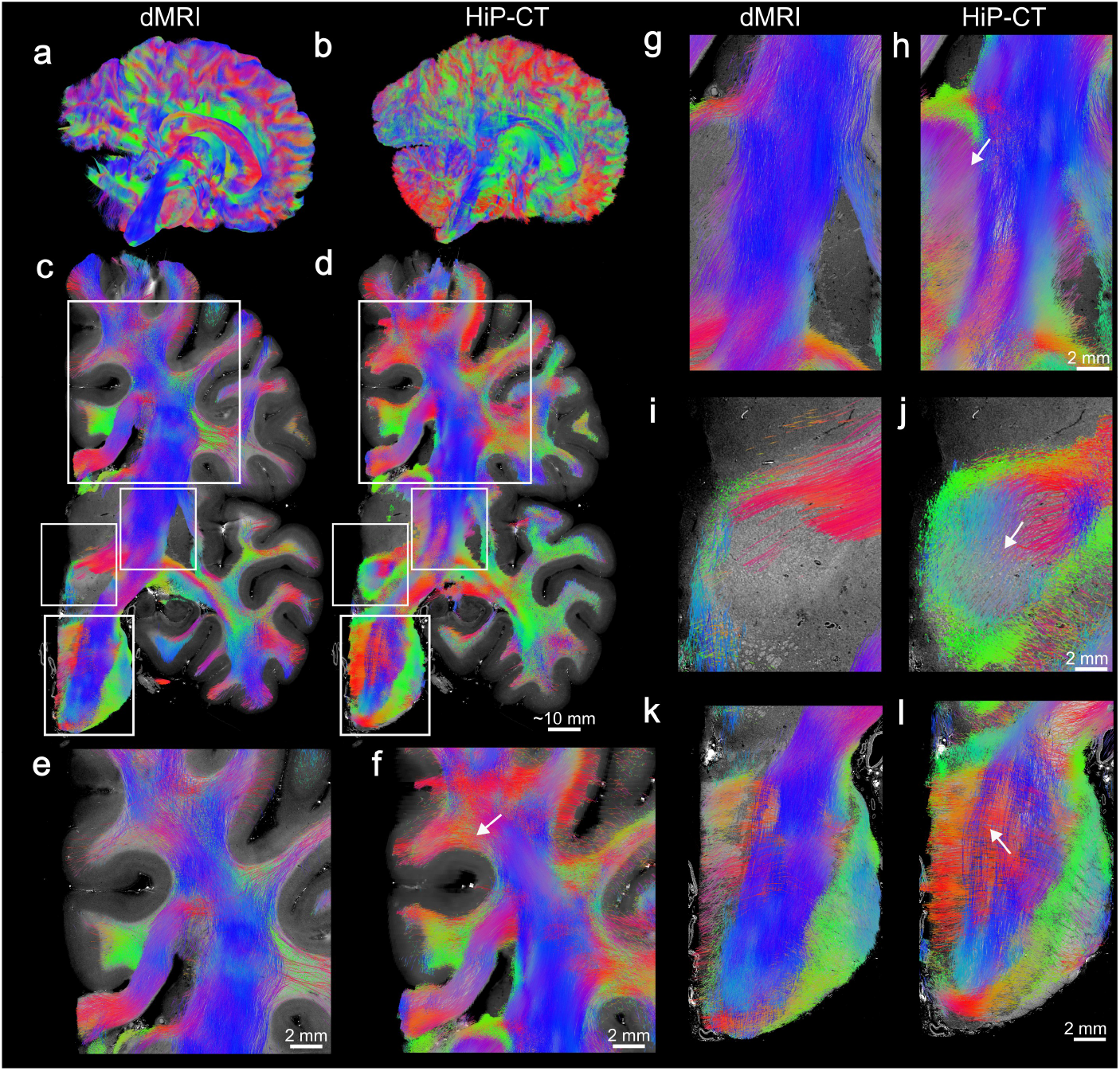
Qualitative comparison of dMRI and HiP-CT tractography at matched 800 µm resolution. Tractograms are generated using Constrained Spherical Deconvolution (CSD) for dMRI (Left columns) and Structure Tensor Analysis (STA) for HiP-CT (Right columns). Streamlines are colored by local orientation (Red: Left-Right, Green: Anterior-Posterior, Blue: Superior-Inferior). (a, b) Whole-brain sagittal 3D views demonstrate high global topological consistency in major white matter architecture. (c, d) HiP-CT coronal slice overlays showing co-registered streamlines in RAS space, with white boxes indicating the regions of interest magnified in the insets below. (e,f) Magnified views of the corona radiata adjacent to the corpus callosum from dMRI (e) and HiP-CT (f). Both methods recover overall dominant orientations, with HiP-CT (f) resolving more complex crossing geometries, including a robust arch of lateral association fibers (top arrow). (g,h) Posterior limb of the internal capsule showing consistent dominant superior–inferior orientation in both modalities, with HiP-CT revealing fibers that peel off the internal capsule and enter the thalamus (arrow). (i, j) Red Nucleus: The dMRI reconstruction (i) shows fibers stopping when they enter the nucleus due to low diffusion anisotropy. In contrast, HiP-CT (j) reveals a dense nexus of fibers that continue to propagate (arrow). (k, l) Pons: Both pontine tractograms highlight orthogonal transverse and longitudinal fiber populations, with HiP-CT capturing transverse fibers threading through the interstitial spaces between distinct longitudinal fascicles.

In the corpus callosum (**Figure 3 e, f**) both methods produce a highly coherent, mediolaterally oriented bundle with smooth streamlines. In the adjacent corona radiata, the overall principal superior-inferior projection and its lateral fanning towards the cortical mantle is broadly consistent between the two modalities. However, HiP-CT tractography captures a broader distribution of orientations (increased green-yellow tracts) suggesting improved sensitivity to fanning and crossing configurations that are partially averaged in dMRI. For instance, the upper arrow in (Figure 3 f) shows a clear emergence of a dense arch of lateral fibers that peel away from the main superior-inferior sheet of fibers towards the cortex. Conversely, this region is sparse and fragmented in the dMRI tractography.

In the internal capsule (**Figure 3 g, h**) both dMRI and HiP-CT tractography recover the principal superior-inferior organization of fibers consistent with the known anatomy of corticospinal and corticopontine projections. HiP-CT (**Figure 3 h)** additionally resolves the transition of fiber bundles into the thalamus which is more challenging to reconstruct with dMRI tractography.

In the red nucleus region (**Figure 3 i, j)**, both dMRI and HiP-CT tractography identify white matter fibers that project into and around the red nucleus. The dMRI reconstruction (**Figure 3 i)** is dominated by laminar, transverse fiber bundles (red) passing superior to the nucleus, effectively rendering the red nucleus itself as a relative tractography void with sparse internal connectivity. In contrast, HiP-CT tractography (Figure 3 j) further traces the interweaving of white matter fibers within the red nucleus that are largely unresolved or under-represented in dMRI tractography.

In the pons (**Figure 3 k, l**), both methodologies capture the macroscopic orthogonal organization, characterized by the crossing descending longitudinal fibers (blue) and transverse fibers (red). While dMRI tractography (Figure 3 k) reconstructs predominantly descending tracts (blue), with transverse fibers (red) less prominent as they traverse laterally through the pons, HiP-CT tractography resolves the interdigitation of transverse and longitudinal fascicles, consistent with the details resolved in the raw HiP-CT images at microscopic resolution **Figure 1**.

Further comparative analysis at a matched mesoscopic resolution (800 µm) demonstrates the capacity of STA-based HiP-CT tractography to resolve complex subcortical white matter architecture that is challenging to reconstruct with CSD-based dMRI tractography. This is particularly pronounced within deep gray matter nuclei, where fiber bundles have small diameters and diffusion anisotropy is low due to partial voluming between gray and white matter [38]. For instance, within the globus pallidus (**Figure 4 a-aii)**), CSD-based dMRI tractography yields a near-complete void of streamlines. In contrast, STA-based HiP-CT tractography, capturing tissue density gradients rather than water diffusion, successfully resolves a dense, multi-directional network of fibers as they traverse the nucleus. Similarly, the fasciculus retroflexus (**Figure 4 b-bii)**) is not resolved in CSD-based dMRI tractography, likely due to its narrow diameter and high curvature. Conversely, STA-based HiP-CT tractography traces the cohesive, curved trajectory of the bundle. Finally, in the highly complex subthalamic region (**Figure 4 c-cii)**), dMRI tractography captures fiber trajectories only partially. HiP-CT tractography, on the other hand, allows a more complete reconstruction of fascicles and nuclei in the region, clearly delineating the zona incerta, the Fields of Forel (H and H2), and the mammillothalamic tract. Notably, however, both methodologies struggle to reconstruct the inferior extent of the mammillothalamic tract. In this lower region, both dMRI and HiP-CT exhibit sparse, fragmented streamlines, reflecting the physical dispersion and loss of coherent bundle density as these fibers approach their terminations in gray matter. Collectively, these comparisons in challenging deep-brain areas demonstrate that, even when STA-based HiP-CT tractography is performed at conventional dMRI resolutions, the structural contrast of HiP-CT facilitates high anatomic fidelity, recovering distinct fascicles that are obscured in dMRI due to the limitations of the diffusion contrast. Further gains in local detail and streamline continuity can be achieved by reducing the supervoxel size, for instance to 400 800 µm (see Supplementary section A.6).

**Fig. 4.**
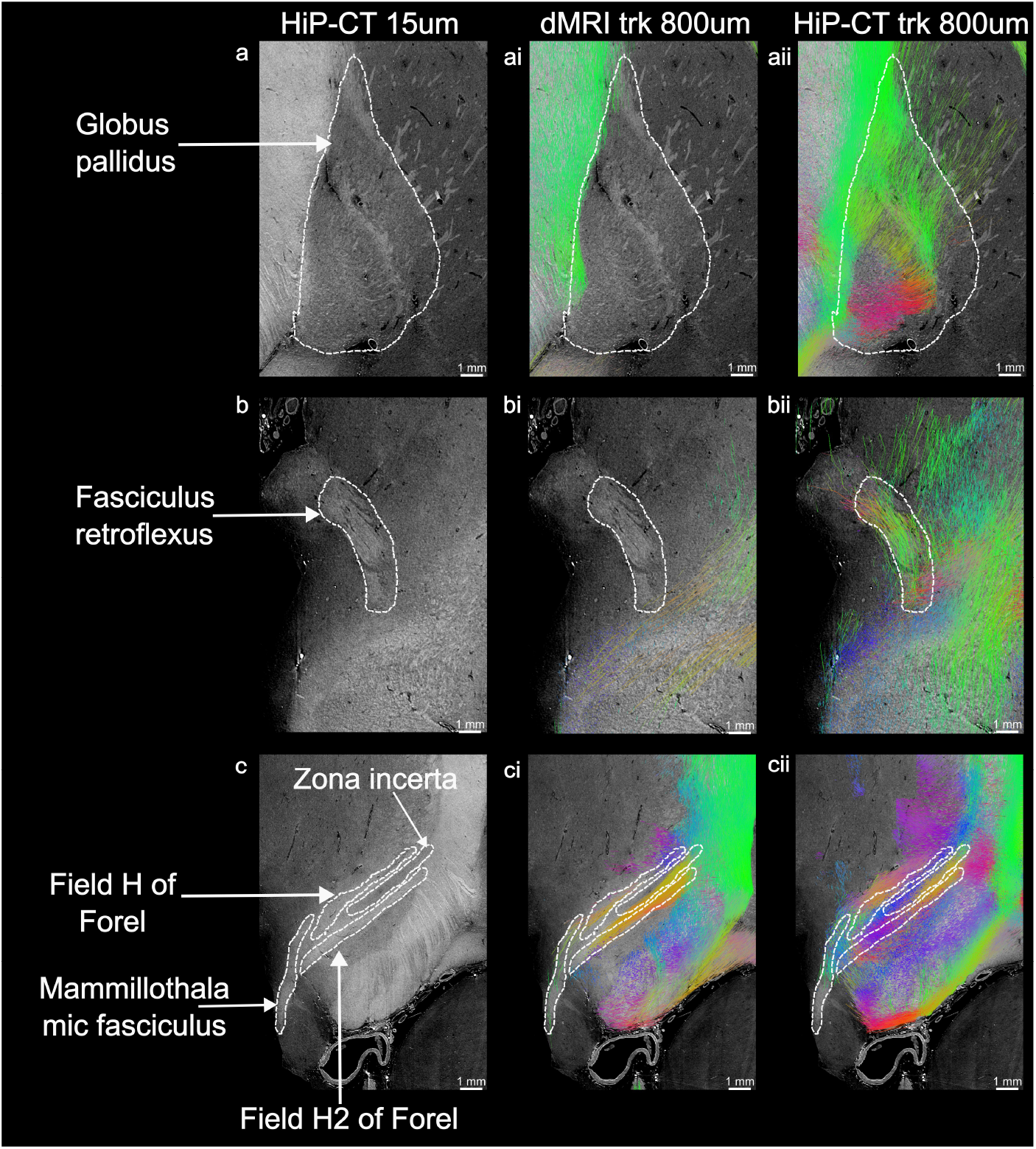
Enhanced resolution of deep brain white matter pathways using HiP-CT structure tensor tractography. Comparison of high-resolution HiP-CT images (left column), CSD-based dMRI tractography (middle column, i), and STA-based HiP-CT tractography (right column, ii). Both tractography reconstructions were evaluated at a matched spatial resolution of 800 µm. White dashed lines outline the anatomical regions of interest in the high-resolution HiP-CT data. (a–aii) Globus pallidus: dMRI tractography (ai) exhibits a near-complete streamline void within this iron-rich deep greymatter nucleus. In contrast, HiP-CT tractography (aii) successfully traces a dense, multi-directional network of traversing fibers. (b–bii) Fasciculus retroflexus: This discrete bundle remains unresolved in dMRI tractography (bi), whereas HiP-CT (bii) robustly recovers its cohesive, curved trajectory. (c–cii) Subthalamic and diencephalic region: In a region of highly complex fiber architecture, dMRI tractography (ci) only partially reconstructs fiber trajectories. Conversely, HiP-CT tractography (cii) allows a more comprehensive reconstruction, clearly delineating distinct sub-structures including the zona incerta, the Fields of Forel (H and H2), and the mammillothalamic tract. Streamlines are colored by local orientation (Red: Left-Right, Green: Anterior-Posterior, Blue: Superior-Inferior).

Together, these results illustrate that, while CSD-based dMRI tractography provides a robust mapping of large-scale white-matter fiber organization, STA-based HiP-CT tractography at a matched resolution can resolve critical additional details. In particular, within transitional zones of white and gay matter (e.g., internal capsule–thalamus interface), deep gray-matter nuclei (globus pallidus), and small-diameter bundles (fasciculus retroflexus, mammillothalamic tract), where dMRI contrast is limited due to partial voluming that leads to low diffusion anisotropy or iron deposition that leads to signal decay, HiP-CT yields denser, more continuous, and anatomically faithful streamlines. This demonstrates the distinct value of HiP-CT as a high-resolution, complementary imaging modality for interpreting and refining dMRI-derived connectivity.

### 2.4 Influence of vasculature on fiber orientations derived from HiP-CT

HiP-CT employs a label-free acquisition in which image contrast arises from variations in electron density, allowing multiple tissue microstructural features with different densities to be identified simultaneously, including vascular networks. **Figure 5 a** shows a segmented vessel network in a 6.54 µm 1024 × 1024 × 1024 Volume Of Interest (VOI) extracted from the pons of the *LADAF-2021* brain scan. At this resolution, it is possible to observe both white matter fiber bundles and blood vessels, their orientation and their spatial interaction. However, this raises the question of whether vascular structures may confound STA-based fiber orientation estimates. To this end, we compare orientation and morphological measurements computed when vasculature is present versus when vasculature has been masked during STA. The masking of vasculature effects is performed using the procedure described in Supplementary section 4.7.

**Fig. 5.**
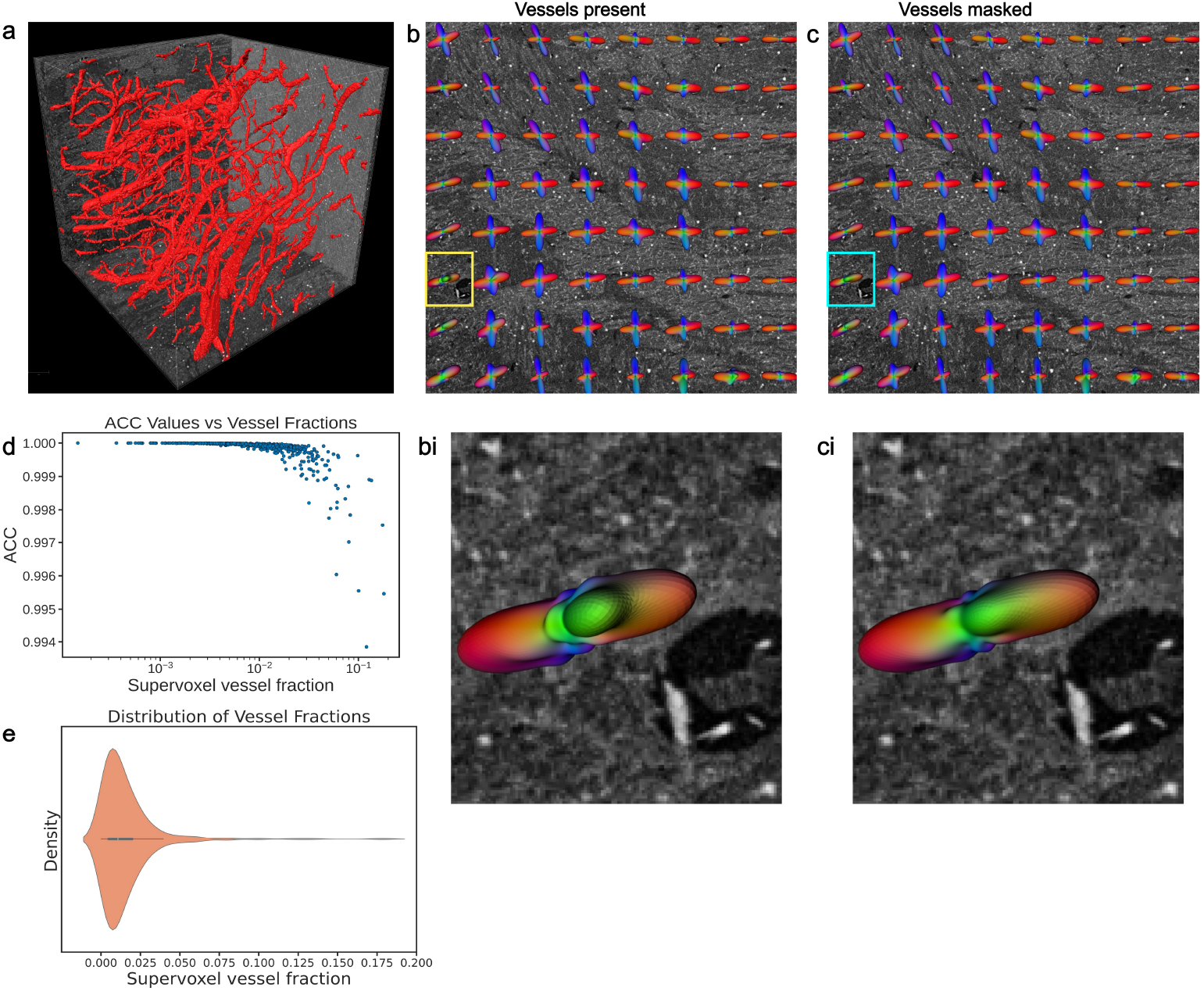
Comparison of STA results with vasculature included and vasculature masked out in a HiP-CT human pons VOI. (a) 3D rendering of the pons VOI showing segmented vasculature (red) overlaid on the HiP-CT data. (b) Coronal slice with overlaid fODF glyphs derived from STA with vessels included. (c) The same slice showing STA results with vessels masked. (bi, ci) Subtle alterations in fODF glyph morphology can be observed when comparing the vessel-included and vessel-masked analyses. (d) ACC between fODFs computed with vessels vs those computed with vessels masked and how that varies with the fraction of vessels within a supervoxel. ACC values generally remain high (∼ 0.99) but exhibit greater variance and a gradual decline as vessel fraction increases. (k) Vessel distribution with the supervoxels. Most supervoxels in the analyzed VOI have a low vessel fraction. Overall, these results indicate a minimal influence of vasculature on STA-derived orientations.

As in previous sections, we compute fODFs within supervoxels of the same size as the dMRI voxel (800 µm). **Figure 5 b, c** shows the dMRI-equivalent supervoxel fODFs computed with vessels present (left) and with vessels masked (right) overlaid on a HiP-CT slice. Visual inspection of fODF glyphs within these supervoxels, consistently shows high similarity in orientation and shape between the vessel-present and vessel-masked conditions. Zoomed insets of supervoxel fODFs that had large discernible vessels are shown in **Figure 5 bi, ci**. Although these paired fODFs all share the same overall orientation whether vessels are present or masked, subtle differences in the shape and symmetry indicate the influence of vessels on the fODFs. The fODF with vessels present (**Figure 5 bi**) exhibits a secondary lobes or minor protrusion along its main axes while the fODF with vessels masked (**Figure 5 ci**) shows less pronounced secondary lobes, with the primary fiber direction appearing more dominant and streamlined. These observations, albeit subtle, indicate that the presence of large vessels can create artifactual fiber directions, while masking them enhances the focus on the true primary fiber orientation.

To account for spatial heterogeneity in vascularization, we investigated whether the vessel volume fraction within a supervoxel modulates their impact on fODF estimation. **Figure 5 d** displays the ACCs between masked and unmasked fODF pairs, plotted against their corresponding supervoxel vessel volume. We observe a monotonic decrease in ACC with increasing vessel volume within the supervoxels. For supervoxels with low vessel volume fractions (∼ 1%), ACC values cluster very close to 1.0, showing minimal orientation divergence. As the fraction of vessel volume increases and approaches (∼ 10%), ACC values generally remain ∼ 0.99 high but exhibit greater variance and a gradual decline. These results indicate that STA-derived fODFs are robust to typical levels of cerebral vasculature ∼ 4% [39]. Voxel-wise anisotropy and shape metrics similarly show highly comparable distributions with and without vessel masking (see Appendix section A.4).

## 3 Discussion

Currently, dMRI is the only technique for imaging the wiring of the brain in vivo. Our results position HiP-CT as a key complementary modality for bridging the critical gap between the mesoscopic resolution of dMRI measurements and the underlying microstructural organization of white matter. We demonstrate that HiP-CT enables non-destructive, isotropic visualization of white matter fibers at the microscale, without the need for contrast agents or destructive sample preparation, overcoming limitations associated with other microscopy techniques that have been used previously to validate dMRI [23, 24, 40, 41]. HiP-CT resolves complex fiber architectures in areas where dMRI is confounded by low diffusion anisotropy. In these challenging areas, HiP-CT allows us to visualize fascicles far smaller than typical dMRI voxels.

Beyond qualitative visualization, we show that STA applied to HiP-CT images enables computation of fiber orientations and tractography in a manner directly analogous to dMRI. Importantly, HiP-CT-derived fODFs recover coherent and crossing fiber configurations within dMRI-sized voxels, paving the way for HiP-CT to be used as a high-resolution reference for evaluating diffusion models. At the level of largescale organization, tractography derived from HiP-CT and dMRI demonstrates strong similarity. Despite relying on fundamentally different physical signal mechanisms, i.e., water diffusion for dMRI and electron density for HiP-CT, both methods reconstruct the major white matter connections consistently. The Direction Encoded Color (DEC) maps for large coherent bundles, such as the corpus callosum and the corticospinal tract, exhibit the same bulk orientations.

Importantly, the complementarity between HiP-CT and dMRI becomes particularly evident in regions where the latter has poor contrast due to low diffusion anisotropy, such as area where white and gray matter intermingle. In transition zones (such as the interface of the internal capsule and thalamus) or within gray matter nuclei (such as the red nucleus and globus pallidus), fiber density decreases, orientation dispersion increases, and MRI signal decays due to high iron content, contributing to poor contrast-to-noise ratio in dMRI. In these settings, dMRI-based tractography is inherently ill-posed, resulting in artifactual truncation of fiber trajctories. HiP-CT, which its derives contrast from electron density can resolve fiber organization in such areas. Our results at the inferior extent of the mammillothalamic tract demonstrate that streamline tractography in both modalities is ultimately confounded by certain complex fiber configurations. We note, however, that tractography parameters for both modalities could be further optimized in such areas, e.g., by relaxing stopping criteria based on the turning angle and/or fODF amplitude or by increasing the maximum stepping angle, to improve propagation in these areas. Such local optimization was beyond the scope of the current study. Furthermore, while this study performed tractography in HiP-CT using fODFs computed the at same resolution as the dMRI data (here 800 µm), the native resolution of HiP-CT (15 µm) allows fODFs to be computed at finer scales. As a proof of concept, we reduced the size of the supervoxel for computing fODFs from HiPCT to 400 µm (see Supplementary section A.6). We found that this preserved the overall reconstructed white-matter organization, while improving the streamline density and continuity of streamlines, demonstrating the scalability of our approach to higher resolutions.

A unique feature of label-free imaging modalities such as HiP-CT is the simultaneous visualization of all soft tissues based on electron density, including white matter, gray matter, and vasculature. This multi-structural contrast enables jointly mapping vascular and axonal architectures within the same intact brain. This opens opportunities for investigating and understanding how vascular topology related to white matter organization, similarly to how high-resolution structural MRI can be used to extract vascular networks for joint analysis with white-matter networks derived from dMRI. While this overall anatomical richness raises the concern that vessels could bias estimates of fiber orientations from STA in HiP-CT, our investigations into this matter revealed that the influence of vessels on orientation and anisotropy metrics is minimal. Even as vessel volume fraction within supervoxels approaches 10%, Angular Correlation Coefficients (ACC) remain high (*>* 0.99). Given that typical cerebral vascular volume fraction is approximately 4% [39], we do not expect vasculature to be a substantial confound in the dominant fiber orientations inferred from HiP-CT data.

Despite these strengths, HiP-CT tractography is subject to several technical limitations. Fundamentally, STA relies on the presence of a sufficient local intensity gradient to accurately capture fiber orientation. In regions exhibiting highly uniform tissue density or weak phase contrast, the lack of a such a gradient renders orientation estimation inherently vulnerable to high-frequency noise. In these areas, minor intensity fluctuations may generate spurious image gradients, leading to instabilities in the tensor eigendecomposition. As observed in our deep-brain comparisons, this can result in noisy or spurious streamlines. These artifacts can be substantially mitigated by anatomically guided masking of such streamlines. Furthermore, the hierarchical image acquisition and reconstruction pipelines used in HiP-CT introduce unique sources of imaging artifacts. For instance, imperfect volume stitching can produce subtle directional discontinuities at scan boundaries, which systematically bias local tensor orientations. Additionally, streak artifacts, common in phase-contrast tomography near sharp density transitions such as trapped air bubbles or fomblin droplets, introduce artificial linear features. These streaks generate strong, erroneous spatial gradients that makes the structure tensor deviate from true directionality in the tissue texture, ultimately resulting in the generation of spurious streamlines (see Supplementary Material section A.5). Collectively, these challenges underscore the necessity for rigorous quality control, advanced denoising algorithms, and targeted artifact correction strategies prior to conducting STA-based tractography in HiP-CT.

Taken together, our results position HiP-CT as a transformative imaging modality for characterizing white matter fiber architectures in a manner that is complementary to dMRI. While dMRI is unique in its ability to probe the human connectome in vivo, HiP-CT provides direct visualization of the underlying tissue microstructure that ultimately drives diffusion contrast. This complementary relationship is particularly powerful in transitional gray-white matter areas and anatomically complex regions around deep brain nuclei, where diffusion anisotropy is low and dMRI tractography terminates prematurely.

Future work will focus on rigorous, quantitative comparisons between co-registered HiP-CT and dMRI-based fODFs and tractography within the same brain. Such investigations will prioritize targeted spatial metrics, including angular concordance, dispersion mapping, and tract continuity analyses. In parallel, the further development of advanced noise-reduction and artifact-mitigation strategies will be essential to improve the robustness of STA-based tractography in HiP-CT. Exploring continuous coordinate-based modeling approaches, such as Implicit Neural Representations, could offer a powerful framework for regularizing these dense orientation fields and smoothing the directional discontinuities caused by volume stitching. Ultimately, these combined efforts will strengthen the integration of HiP-CT and dMRI, towards true multi-modal modeling of white matter microstructure and macroanatomy in health and disease.

## 4 Methods

### 4.1 Sample collection and preparation

Two human brain samples were scanned and analyzed within the scope of this work. The first sample was a left cerebral hemisphere (donor ID: I58) obtained from an 81-year-old male donor through the Massachusetts General Hospital (MGH) Autopsy Suite. Consent for body donation and use in research was obtained in accordance with institutional and national regulations. The post-mortem interval was less than six hours. The I58 specimen was initially fixed in 10% formalin for at least 2 months then stored in 2% Periodate-Lysine-Paraformaldehyde (PLP). After dMRI scanning, the specimen was transported to the European Synchrotron Radiation Facility (ESRF, Grenoble, France) under the approval of the French Ministry of Health for import and imaging of human tissues. The second sample was a whole brain from a 63-year-old male body donated to the Laboratoire d’Anatomie des Alpes Françaises (LADAF) following the current French legislation for body donation. Written informed consent was obtained antemortem and all dissections respected the memory of the deceased. Transport and imaging protocols were approved by the French Health Ministry.

Prior to HiP-CT imaging, both samples were immersed in a 4% neutral buffered formalin bath for 4 days, then partially dehydrated progressively until 70% ethanol with successive baths. To prevent bubble formation during scanning, which would create artifacts and reduce image quality, the samples underwent in-line degassing. The samples were mounted in a container filled with agar-ethanol gel to maintain the brain in position during scanning. Finally the container was sealed until imaging acquisition. For more information on the sample preparation protocol, see Brunet et al. [42].

### 4.2 Diffusion MRI acquisition and analysis

Prior to HiP-CT data acquisition, dMRI data for the I58 hemisphere were acquired at the MGH Martinos Center on the 3T Connectome 1.0 scanner (maximum gradient strength = 300 mT/m) [9], using a custom-designed 48-channel receiver *ex vivo* coil [43] and a 3D segmented EPI diffusion-weighted sequence optimized for *ex vivo* dMRI [44]. Diffusion data were acquired at 800*µm* isotropic resolution across 32 diffusion-encoding directions with *b* = 4000 *s/mm*^2^ and 40 directions with *b* = 10000. Eight non diffusion-weighted volumes (*b* = 0) were also acquired. Additional acquisition parameters include: echo time TE = 74 ms, repetition time TR = 500 ms, anterior-posterio phase-encode direction, number of signal averages = 1, number of segments = 5. Prior to MRI scanning, the brain was transferred into a sealed plastic bag filled with Periodate-Lysine-Paraformaldehyde (PLP) and checked daily to ensure elimination of most air bubbles that would cause susceptibility artifacts. Image pre-processing steps included denoising [45], correction of image distortions due to B0 inhomogeneities and eddy currents [46, 47], along with correction of distortions due to gradient non-linearity. The warp fields estimated for the susceptibility, eddy-current, and gradient nonlinearity distortions were combined and applied to each image volume in a one-step resampling process to minimize smoothing effects due to interpolation. Images were then corrected for intensity biases and fiber Orientation Distributions Functions (fODFs) were reconstructed using a multi-shell, multi-tissue constrained spherical deconvolution approach (MSMT-CSD) in MRtrix3 [48–50]. The fODFs were used to propagate probabilistic tractography from 5 random seed locations in every white-matter voxel [51].

### 4.3 HiP-CT acquisition

All HiP-CT imaging was performed at the European Synchrotron Radiation Facility (ESRF) on beamline BM18, which is optimized for propagation-based phase-contrast tomography using the Extremely Brilliant Source (EBS). Acquisition protocols followed previously published HiP-CT procedures [3, 42]. No contrast agent was required. A parallel polychromatic X-ray beam was filtered with different combinations of sapphire, SiO_2_, and glassy carbon attenuators, providing average energies between 90 and 120 keV depending on the scan. The propagation distance was set to 15 m for full-field overview scans of the whole organ and between

#### Donor I58

An overview scan of the entire left hemisphere was performed at an isotropic voxel size of 15 µm. The acquisition used quarter-acquisition mode [3] with a z-series of 32 overlapping tomograms, each covering 5 mm in the vertical direction with 30% overlap. Each tomogram consisted of 15,000 projections over 360^°^. The complete acquisition required approximately 20 hours. A subsequent local zoom of the red nucleus was acquired at 6.21 µm isotropic voxel size. This dataset was acquired in half-acquisition mode with a z-series of 7 tomograms, each separated by 5 mm. Each tomogram consisted of 15,000 projections. The propagation distance was reduced to 5 m.

#### Donor LADAF-2021-17

For this donated brain sample, a high-resolution local zoom was performed at an isotropic voxel size of 6.54 µm. The scan was carried out in half-acquisition mode with 6000 projections over 360^°^. The mean beam energy was 120 keV and the propagation distance was 10 m. The lateral field of view was approximately 25 mm.

### 4.4 Cross-modal image registration

To directly compare fiber orientation estimates from dMRI and HiP-CT, we developed a multi-scale, cross-modal, registration pipeline using the dMRI scan 800 µm as the moving image and the HiP-CT scan 15 µm as the fixed image. The HiP-CT volume was first downsampled from 15 µm to 240 µm to bridge the resolution disparity with the 800 µm dMRI. This served as a low-pass filter that harmonized macroscopic anatomical features between the two scans and significantly reduced both the memory footprint and the degrees of freedom in the estimation of dense non-linear deformation fields. The average *b* = 10000 volume, normalized by the average *b*_0_ volume was used as the moving image, providing a contrast similar to that of the HiP-CT image.

We utilized a pipeline consisting of rigid, affine and deformable image registration, implemented in ITK-SNAP [52] and ANTsPy [53]. Registration was initialized using a manually created rigid transformation in ITK-SNAP to resolve gross orientation differences. Optimization was then performed using a hybrid metric strategy: an initial affine stage driven by mutual information (32 bins) to correct global scaling and shearing, followed by a symmetric normalization [54] deformable stage driven by neighborhood cross-correlation (radius = 4 voxels). Finally, whole-hemisphere dMRI tractography streamlines were mapped from native dMRI space to HiP-CT space by applying the inverse affine and inverse warp fields to the streamline coordinates.

### 4.5 HiP-CT analysis

#### Structure Tensor Analysis

Voxel-wise fiber orientations were estimated using an image processing technique known as *structure tensor analysis*, which has been previously applied for this purpose in numerous microscopy modalities, including polarized light imaging, polarization-sensitive optical coherence tomography, light-sheet microscopy and synchrotron micro-CT datasets [22, 24, 27, 32, 40, 55–58]. The structure tensor is the gradient of image intensities in a local neighborhood. After smoothing image intensities with a Gaussian kernel of standard deviation *σ*, K_*σ*_ i.e *I*_*σ*_ = K_*σ*_ ∗ *I*, and the structure tensor is obtained by computing the partial derivatives in the *x, y, z* dimension. More efficiently, this can be implemented as a convolution of the image intensity with the derivative of a Gaussian. Next, a gradient square tensor is calculated for each point in the image by taking the dyadic product of the resulting gradient vector with itself:

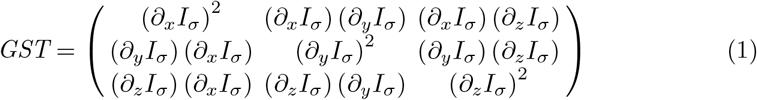

Finally, aggregation of tensor information within a local neighborhood is done by filtering the gradient square tensor with a Gaussian filter with standard deviation *ρ* resulting in the structure tensor **S**_*ρ*_:

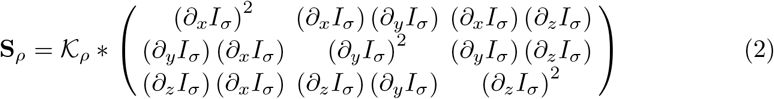

In essence, *σ* is used for noise reduction and setting the scale of features considered by the gradient, while *ρ* is used for spatial averaging of features over a neighborhood to get a stable estimate of orientations. Structure tensor analysis was performed using the structure-tensor Python package [59] provided at: https://github.com/Skielex/structure-tensor. Gaussian kernels with a noise scale (*σ* = 2.0) and integration scale (*ρ* = 4.0) were used in this analysis. The eigendecomposition of the structure tensor is performed to obtain three eigenvalues, *λ*_1_ ≥ *λ*_2_ ≥ *λ*_3_, and their corresponding orthogonal eigenvectors, **v**_1_, **v**_2_, **v**_3_. The eigenvector *v*_3_ corresponding to the smallest eigenvalue represents the direction of minimal local variation in image intensity, here assumed to represent fiber orientation[60]. Various metrics have been proposed in the literature to compute anisotropy from the eigenvalues[32, 40, 60–62]. Highly anisotropic, fiber-like image texture is indicated by a high value of

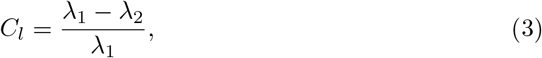

while plane-like structures are described by a high value of

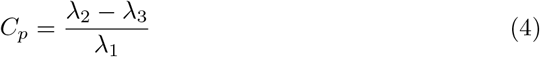

and isotropic structures are described by a high value of the spherical shape measure

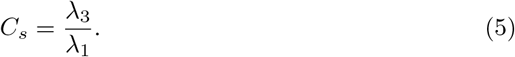

In a manner analogous to fractional anisotropy (FA) from diffusion tensor imaging, a scalar ranging from zero to one that quantifies the directional coherence, is calculated as follows:

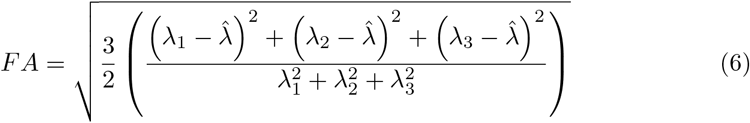

where 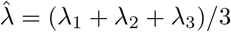 is the mean of the three eigenvalues.

Given the massive scale of the high-resolution HiP-CT data (∼ 190GB for the 15 µm whole brain hemisphere), applying continuous 3D convolutions in memory is computationally intractable for standard research workstations. To ensure our methodology remains accessible and reproducible for the broader neuroscience community, we developed a highly scalable, open-source computational pipeline based on out-of-core processing. The raw tomographic data was first converted into the Zarr format (https://github.com/zarr-developers/zarr-python), enabling compressed, chunk-based data storage and rapid spatial querying. To prevent the edge artifacts inherent to filtering chunked boundaries, each spatial chunk was dynamically loaded into memory with an overlapping padding margin, determined by the filter truncation and the maximum standard deviation of the Gaussian kernels. Structure tensor computations on these chunks were distributed across multiple CPU threads using the Dask parallel computing library (https://github.com/dask/dask). Following computation, this padding was systematically discarded before performing the eigen-decomposition. Finally, to manage the massive data expansion resulting from the tensor outputs, the computed dense fields of eigenvalues and eigenvectors were serialized directly back to disk as Zarr arrays. By utilizing this chunked, lazy-loading architecture, we ensure that the multi-terabyte directional fields remain highly accessible for seamless downstream quantification (such as estimating fODFs), allowing researchers to rigorously analyze the data while strictly adhering to conventional hardware memory bounds.

#### Estimation of fODFs from the HiP-CT structure tensor

The fODF is a formalism widely used in dMRI to model multiple fiber orientations per voxels for tractography[63, 64]. Here we also compute fODFs from the HiP-CT structure tensor, to allow direct comparison of fiber orientation estimates derived from dMRI and HiP-CT.

HiP-CT derived fODFs are obtained using the high-resolution map of the structure tensor eigenvector **v**_3_. First, the high-resolution vector field (15 µm) is partitioned into groups of super-voxels equivalent to the voxel size of the dMRI data (800 µm).

Next, within each supervoxel, the vectors are aggregated into a spherical histogram parametrized by azimuth *ϕ* ∈ (0, 2*π*) and elevation angles *θ* ∈ (0, *π*). This yields a discretized empirical distribution of orientations on the unit sphere. The resulting histograms were then fitted to eighth-order (*l*_*max*_ = 8) spherical harmonic functions as per **Equation 7** to obtain a continuous probability distribution function representation in each supervoxel.

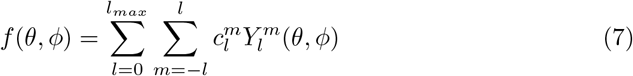

The HiP-CT–derived fODFs are visualized as 3D glyphs in the same manner commonly used for dMRI-based fODF depiction in the literature. Spherical harmonic estimation and glyph rendering are implemented in Python using code adapted from established dMRI analysis libraries, including DIPY [65] and Fury [66].

This methodological approach is consistent with previous studies that have used structure-tensor–derived fODFs to validate dMRI against histology, polarized light imaging, and synchrotron micro-CT [22, 24, 32, 41, 67].

While metrics derived from the eigenvalues, such as FA, can quantify the degree of orientation preference, they tend to decrease in regions where multiple orientations coexist because the structure tensor becomes more isotropic. To overcome this ambiguity, metrics derived from the fODF of the structure tensor can provide complementary information. Among these, Generalized Fractional Anisotropy (GFA) is particularly suitable as it quantifies the angular variance of the ODF amplitudes on a sphere and is calculated as:

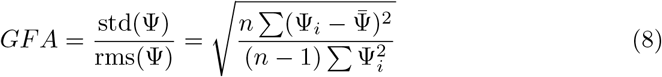

where Ψ_*i*_ are the sampled fODF values on the sphere, *n* is the number of samples (directions) per ODF, and 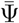 is the mean of the fODF.

While structure-tensor FA decreases in regions with multiple fiber orientations, the fODF often retains multiple distinct peaks, leading to relatively high GFA values in the same region. Given this robustness, we use the GFA in the stopping criterion for tractography.

As in the previous section, this part of the pipeline was also designed around open-source tools and out-of-core processing. This allowed the massive spatial aggregation of billions of high-resolution eigenvectors into 800 µm supervoxel fODFs to be efficiently parallelized across standard multi-core CPUs without exceeding conventional memory limits or CPU cluster if available. The resulting SH coefficients were subsequently serialized back to disk as compressed Zarr arrays, ensuring that the resulting fODF fields are lightweight, easily distributable, and seamlessly compatible with standard open-source tractography libraries like DIPY [65].

### 4.6 Structure tensor based tractography

We applied probabilistic tractography, as implemented in the DIPY library [65], to the fODFs derived from the HiP-CT structure tensors.

#### Seeding

Streamlines were seeded in voxels where the structure-tensor derived FA exceeded 0.42. This empirical threshold was selected based on the voxel-wise FA distribution, where it approximately corresponded to the median FA and optimally differentiated the highly coherent, anisotropic structure of the white matter from the more isotropic surrounding gray matter.

#### Stopping criterion

Streamlines were terminated when they entered voxels with a GFA value below 0.7, preventing propagation into isotropic or noise-dominated regions. GFA was used because it remains high even in crossing regions, hence fiber tracing can continue even through complex white matter configurations.

#### Propagation

Streamlines were propagated by sampling from the structure tensor-derived fODFs at each step. We empirically chose the propagation step size to be a quarter of the dMRI voxel size, and the maximum turning angle between steps to be 30^°^.

### 4.7 3D vascular segmentation

The image contrast in HiP-CT is non-specific, capturing a variety of brain structures that include cell clusters and blood vessels. In HiP-CT imaging, vessels appear as tubular structures of high or low intensity, depending on the presence of residual blood in the vessel during sample preparation. Vascular structures have been identified in HiP-CT data spanning from large vessels (cm in diameter) down to artierioles/venuoles (10’s of *µm* [68, 69]). When vasculature is present, it can contribute to strong linear structures or regions that form sheet-like structures or isotropic regions, as demonstrated by [60]. As vessels often follow similar trajectories as fibers, it is important to determine whether the presence of vasculature could act as a confounding factor in structure tensor analysis of HiP-CT imaging data, artificially dominating the estimated anisotropy and principal orientations in white matter.

To investigate this we sought to compare the results of structure tensor analysis performed on HiP-CT data both *with* and *without* the inclusion of vascular structures. The Volume Of Interest (VOI) used in this study was a 6.54 µm 1024 × 1024 × 1024 volume extracted from the pons of the *LADAF-2021* brain scan. To perform accurate segmentation of vasculature, we adopt a human-in-the-loop segmentation framework. This approach leverages the strengths of both a convolutional neural network (CNN) for automated segmentation and human expert review in a collaborative process designed to achieve high segmentation accuracy for vascular structures while minimizing the extensive manual annotation typically required for large, complex datasets. To achieve this desired objective, the deep learning interface, “Segmentation Wizard”, developed by ORS (Object Research Systems) Dragonfly Inc [70] was used. Dragonfly takes Keras-based Python-encoded Convolutional Neural Networks and provides an interface that facilitates parameter tuning, iterative training, and inference of deep learning models on new inputs [70]. At a high-level, the workflow adopted in this study takes the following schema:

1. An initial set of high-quality training slices from the VOI are created by manual annotation.
2. A suitable deep learning model architecture is chosen.
3. The model is trained and inference made on the rest of the volume.
4. Results are inspected, regions of poor performance are corrected and added to the training data, and model hyperparameters adjusted if needed.
5. Step 3 and step 4 are repeated until qualitative and quantitative user-defined metrics are met.

This workflow is summarized as shown in **Figure A1**.

Within the ORS Dragonfly Segmentation Wizard, the segmentation process began by selecting a slice that best represented the full diversity of vasculature features. A segmentation frame was then defined encompassing the entire slice, and manual annotation of vasculature and background was performed using a local Otsu [71] round brush tool, yielding 7 initial exemplary training slices. A 2.5D pretrained U-Net was chosen for semantic segmentation with details of the model architecture, training parameters, and data augmentation outlined in **Table A2**. Following the first training round, we performed inference across the full volume and evaluated the resultant segmented vascular network using metrics such as Dice score, connected component count, labeled voxel count, contribution of the largest connected component, and total labeled voxel volume. Although the model adeptly identified small vessels, it struggled with larger ones, prompting us to annotate eight additional slices, bringing the total to 15, and retrain the model. Further iterations with 22, 33, and 53 slices enriched the training set with diverse vascular characteristics, aiming to enhance generalization. We halted training when quantitative metrics indicated diminishing improvements in the vascular network. **Figure A2** shows the change in the segmented network’s characteristics after each training run.

As the model undergoes iterative training and refinement from the first to the fourth iteration, metrics such as labeled voxel count, labeled voxel volume, largest connected component contribution, and vessel length increased, reflecting improved segmentation, before declining at the fifth iteration. These trends aligned with validation patch performance, as shown in **Table A3**. Ultimately, the fourth iteration model delivered the strongest balance of quantitative performance (validation dice = 0.9264) and qualitative anatomically plausible vascular networks (see **Figure A3**). Consequently, this model was saved and picked as the best candidate to perform segmentation of vasculature in HiP-CT brain images. This approach minimized manual annotation effort while ensuring robust vascular segmentation for subsequent analysis of vasculature’s impact on white matter structure tensor analysis.

#### Masking out vasculature effects

To investigate the impact of vasculature on white matter measurements, we investigated two approaches to reduce vascular contribution:

##### 1. Pre-Structure Tensor Vessel Masking

In this approach, segmented vascular structures were removed from the Volume of Interest (VOI) before computing the structure tensor. Vasculature was masked out from the VOI by matrix multiplication with an inverted vessel mask, followed by computation of the 3D structure tensor and eigen decomposition to obtain eigenvalues and eigenvectors.

##### 2. Gradient-Level Vessel Masking

This approach aimed to suppress vascular signals at an intermediate stage of the structure tensor computation. The original VOI, containing both white matter fibers and vasculature, was first convolved with the derivative of a Gaussian filter with standard deviation *σ* to compute image intensity gradients. Next, vasculature gradients were suppressed by applying an inverted vessel mask directly to the intensity gradients. Finally, tensor information was aggregated within a local neighborhood defined by a second Gaussian filter (standard deviation *ρ*) before performing eigen decomposition to obtain eigenvalues and eigenvectors.

The effectiveness of each masking approach was evaluated based on its ability to minimize the gradients associated with vasculature. Gradients reflect the rate of intensity change within the volume. Therefore, to exclude the contributions of vasculature, the aim is to make the intensity gradients generated by vessels as minimal as possible. As evidenced in **Figure A4**, the pre-structure tensor vessel masking approach generated artificially high gradients at the vessel regions, while the gradient-level masking approach effectively suppressed vascular gradients, producing a structure tensor that was minimally influenced by the presence of vessels. This observation shows that pre-masking the VOI creates voids in intensity, leading to artificially high gradients in vessel regions.

Based on this evaluation, the gradient-level vessel masking approach was selected for all subsequent structure tensor analyses aimed at investigating the influence of vasculature on characterizing white matter fiber properties.

#### Investigating the effects of vessel masking

To quantitatively assess the impact of vessel masking on HiP-CT scalar anisotropy metrics (such as FA), we conducted paired t-tests of the metrics within the dMRI-equivalent supervoxels. For each supervoxel, the mean value of the metric was calculated before vessel masking and after vessel masking. Paired t-tests were performed on these corresponding means to investigate any statistically significant difference. The effect size, which quantifies the magnitude of the difference and practical significance was calculated using Cohen’s d. Additionally, as a non-parametric alternative and robustness check, we performed a Wilcoxon signed-rank test on the paired differences of the mean anisotropy values across the super-voxels. This test evaluates whether there is a significant difference between the paired differences of observations and provides a robust measure of effect even when the distribution of differences deviates from normality.

To further evaluate the effect of vessels on fiber orientation, we computed the Angular Correlation Coefficient (ACC) between fODFs computed with vessels present and fODFs computed when vessels were masked. ACC provides an estimate of how closely a pair of fODFs are related on a scale of -1 to 1, using their spherical harmonic (SH) coefficients *u* and *v*:

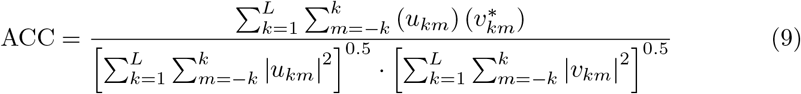

## 5 Acknowledgment

We sincerely thank the donors and their families for their invaluable contributions. Results from such research can potentially increase humanity’s overall knowledge that can then improve patient care. Therefore, these donors and their families deserve our highest gratitude. We gratefully acknowledge ESRF beamtimes md1290 and md1389 on BM18 as sources of the data.

## Ethics approval and consent to participate

Samples were obtained via Massachusetts General Hospital, Harvard Medical School body donation program. Consent for body donation and use in research was obtained in accordance with institutional and national regulations. Specimens were transported to the European Synchrotron Radiation Facility (ESRF, Grenoble, France) under the approval of the French Ministry of Health for import and imaging of human tissues.

## Consent for publication

All authors have reviewed the manuscript and consent to publication.

## Data availability

Public HiP-CT datasets are available through the Human Organ Atlas Hub, hosted by the European Synchrotron Radiation Facility (ESRF) (https://human-organ-atlas.esrf.fr/). Both the I58 dataset and the LADAF-2021-17 are available for browsing and download via this portal.

## Materials availability

Not applicable.

## Code availability

Custom code, scripts, and pipelines utilized for the HiP-CT Structure Tensor Analysis (STA) and corresponding tractography are open-source and freely available on GitHub at https://github.com/R-icntay/towards_dmri_hipct. The human-in-the-loop vascular segmentation was performed using the “Segmentation Wizard” module within Dragonfly software (Object Research Systems, Inc., Montreal, Canada). Dragonfly is commercially available, but free licenses are available for non-commercial academic research.

## Author contribution

Author contribution (according to the [CRediT](https://credit.niso.org/) taxonomy):

*Conceptualization* E.W., M.C., P.D.L., C.L.W.

*Data curation* E.W., C.M., J.B., S.Y.H, P.T.

*Formal analysis* E.W., C.L.W.

*Funding acquisition* A.Y, P.D.L., C.L.W.

*Investigation* E.W., M.C., C.M.

*Methodology* E.W., C.M., M.C., C.L.W

*Project administration* A.Y., P.D.L., C.L.W.

*Resources* S.Y.H., J.B.

*Software* E.W., Y.B,

*Supervision* M.C., A.Y., P.D.L., C.L.W.

*Validation* M.C., C.M., A.K., B.F, A.Y., P.D.L., C.L.W.

*Visualization* E.W., M.C., A.K., A.S.

*Writing – original draft* E.W., M.C., C.L.W

*Writing – review & editing* M.C., C.M., Y.B., J.B., B.F, A.Y., P.D.L, C.L.W

## Appendix A Supplementary Information

### A.1 HiP-CT Acquisition Parameters

Table A1 details the specific synchrotron X-ray phase-contrast imaging setups used for the hierarchical scanning of the brain samples. Because HiP-CT relies on a propagation-based phase-contrast setup, parameters were specifically optimized for each target resolution. Data for the macroscopic overview scan (Sample I58) was acquired at a 15.13 µm voxel size using an energy of 90 keV, requiring 32 vertical scans to capture the extensive 142.42 mm lateral field of view. In contrast, the high-resolution, localized volume (Sample LADAF-2021-17) was acquired at a 6.54 µm voxel size at 120 keV, utilizing a shorter propagation distance (10 m) and a tighter 25.15 mm field of view. Both acquisitions utilized multiple vertical steps and optimized attenuators to manage beam flux and minimize artifacts across the sample volumes.

**Fig. A1.**
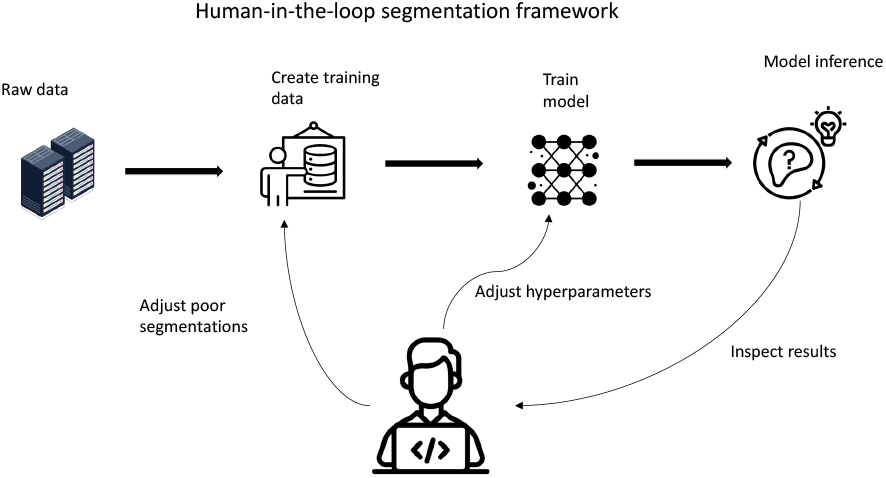
Human-in-the-loop (HITL) workflow for multi-scale vascular segmentation in HiP-CT data. The iterative process leverages a 2.5D U-Net within the ORS Dragonfly Segmentation Wizard. Initial manual annotations of a 6.54 µm VOI from the pons establish a baseline model. Subsequent cycles of full-volume inference, human expert review, targeted error correction, and training set expansion progressively refine the model’s ability to isolate complex, multi-scale vascular networks from the surrounding structural anatomy.

**Table A1.** HiP-CT scanning parameters.

| Sample ID | Voxel ($\mu\text{m}$) | Prop. dist. (m) | Attenuator | Energy (keV) | Lat. FOV (mm) | Proj. | Expo (ms) | Accum. | Acq. mode | Scan time (s) | N scans | Vert. FOV (mm) | Vert. step (mm) |
| --- | --- | --- | --- | --- | --- | --- | --- | --- | --- | --- | --- | --- | --- |
| I58 | 15.13 | 15 | Sapphire 10mm, SiO<sub>2</sub> 15mm block | 90 | 142.42 | 15000 x 2 | 14 | 5 | quarter | 1071 | 32 | 6.05 | 5 |
| LADAF-2021-17 | 6.54 | 10 | Mo 0.21mm | 120 | 25.15 | 6000 | 60 | 4 | half | 369 | 9 | 13.4 | 7 |

### A.2 Human-in-the-loop Vessel Segmentation

As noted in the main text, HiP-CT is a non-targeted imaging modality that naturally resolves a variety of structural features alongside white matter, most prominently dense vascular networks spanning from large vessels down to micro vasculature [68, 69]. Because these vascular structures exhibit strong tubular or sheet-like geometries, they generate pronounced spatial gradients in the phase-contrast images [60]. During structure tensor analysis, these non-axonal gradients can artificially inflate local eigenvalues, thereby introducing spurious anisotropy and biasing the orientation measurements intended to characterize white matter architecture.

**Fig. A2.**
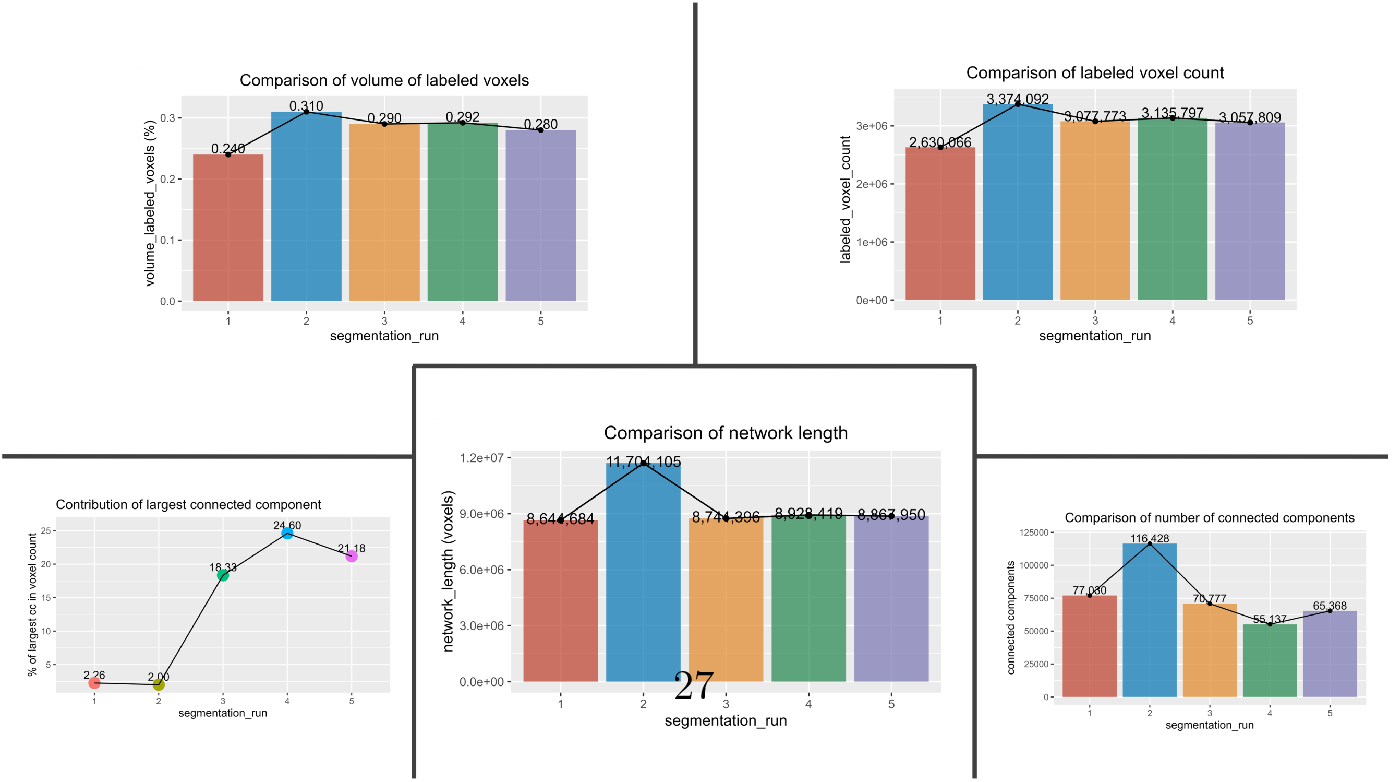
Evolution of quantitative vascular segmentation metrics across iterative training rounds. The graph tracks key performance indicators—including validation Dice score, connected component count, and total labeled voxel volume—as the human-annotated training set expanded from 7 to 53 slices. Performance metrics peaked at the fourth training iteration (33 slices) before declining at the fifth iteration, indicating the onset of diminishing returns and model overfitting.

**Fig. A3.**
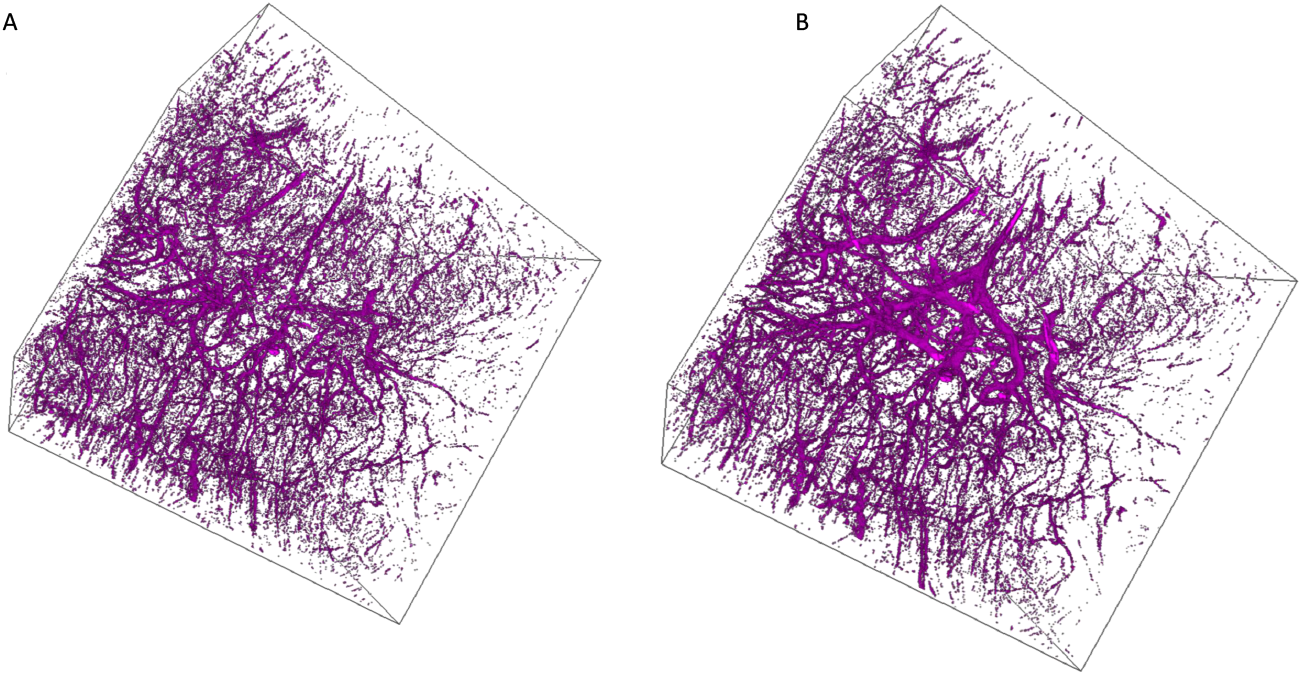
Qualitative improvement in multi-scale vascular network segmentation. **(a)** The segmented vessel network following the initial training iteration captures prominent large vessels but suffers from severe structural fragmentation and fails to resolve microvasculature detail. **(b)** The optimized network from the fourth training iteration demonstrates significantly improved anatomical plausibility. It successfully resolves both massive tubular vessels and the dense, continuous microvascular bed.

**Table A2.** Convolutional Neural Network architecture and training hyperparameters. Parameters define the 2.5D U-Net deployed for the semantic segmentation of vasculature. The architecture utilizes a depth of 6 and 3-slice input dimensions to capture continuous 3D spatial context, while extensive data augmentation (rotations, flips, zooming, and shearing) was applied to force model generalization across the diverse morphological scales of the vascular network.

| Model Architecture | Training Parameters | Data Augmentation |
| --- | --- | --- |
| Depth level: 6 | Batch size: 512 | Horizontal/vertical flips |
| Initial filter count: 64 | Patch size: [64, 64, 1] | Rotation: 180.0° |
| Input dimension: 2.5D, 3 slices | Stride ratio: 0.25<br>Loss function: Binary cross-entropy | Zoom: [0.9, 1.1] |
|  | Optimization: Adadelata<br>Epochs: 100 | Shear: 2.0 |

**Table A3.** Summary of validation performance per HITL training iteration. Validation Dice scores were recorded after 100 epochs for each of the five training rounds. The model from Round 4 achieved the highest quantitative accuracy (Dice = 0.9264) and was subsequently selected to generate the final vascular masks for the structure tensor analysis pipeline.

| Training Round | Epochs Trained | Validation Dice |
| --- | --- | --- |
| 1 | 100 | 0.8325 |
| 2 | 100 | 0.7986 |
| 3 | 100 | 0.8778 |
| 4 | 100 | 0.9264 |
| 5 | 100 | 0.9186 |

To quantitatively assess and control for this confounding factor, we developed a pipeline to segment and mask the vasculature prior to tractography. This section details the human-in-the-loop (HITL) deep learning framework utilized to isolate these networks. By combining the high-throughput inference capabilities of a Convolutional Neural Network (CNN) with iterative human expert review, we achieved accurate, multi-scale vascular segmentation while minimizing the prohibitive manual annotation typically required for massive tomographic datasets.

### A.3 Masking out vasculature effects

As detailed in the main text, safely removing the confounding influence of vasculature from the structure tensor analysis requires careful timing within the computational pipeline. Because the structure tensor calculates orientation based on spatial intensity derivatives, simply zeroing out the vessels in the raw image volume (Pre-Structure Tensor Masking) creates sharp, artificial boundaries between the intact tissue and the empty vessel voids. The derivative of this artificial step-edge manifests as a massive, spurious gradient. To visualize why this approach is detrimental, this section demonstrates the artificial edge effects caused by pre-masking compared to the clean signal suppression achieved by applying the mask to the spatial derivatives directly (Gradient-Level Masking).

**Fig. A4.**
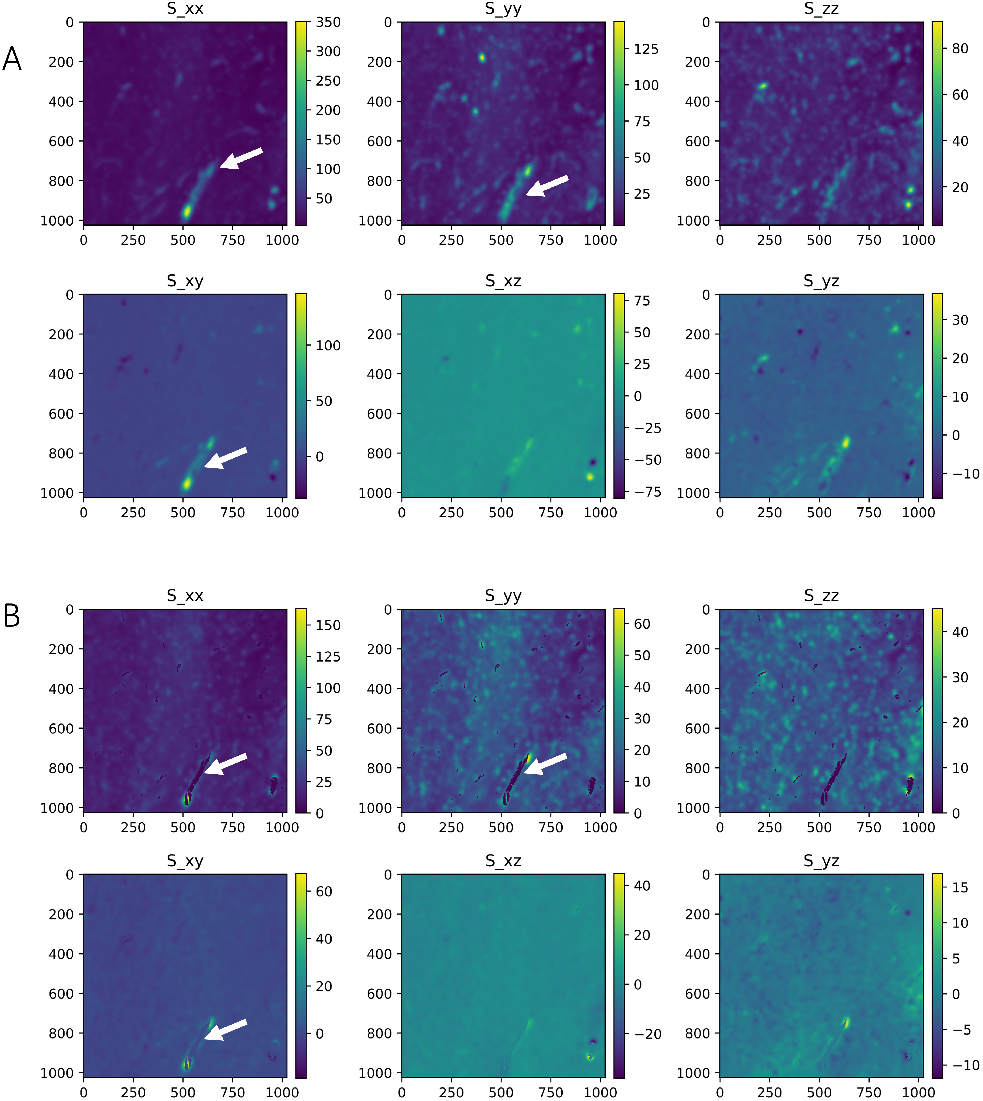
Comparison of vasculature masking strategies on structure tensor gradient computation. **(A)** Pre-Structure Tensor Masking: Removing vessels directly from the raw Volume of Interest (VOI) creates sharp artificial boundaries, inadvertently generating massive, spurious gradients (high intensity) around the vessel voids. **(B)** Gradient-Level Masking: Applying the mask after computing the spatial derivatives effectively suppresses the vascular signal, resulting in the intended low-gradient regions without introducing boundary artifacts.

### A.4 Effects of vasculature on white matter fiber anisotropy

We extended vasculature effect analysis to quantify the specific effects of vasculature on white matter anisotropy and shape metrics, namely Fractional Anisotropy (FA), linearity (fiber-like symmetry), and planarity (plane-like symmetry). Voxel-wise comparisons of these metrics, computed both with and without gradient-level vascular masking, are presented as violin plots in **Figure A5 a, b, c**.

The violin plots reveal highly similar and largely symmetric distributions across both conditions, exhibiting comparable interquartile ranges for FA, linearity, and planarity. This visual similarity suggests that the overall population characteristics of these shape metrics within white matter voxels remain largely unperturbed by the presence of unmasked vessels. While statistical testing (Wilcoxon signed-rank and paired t-tests) yielded statistically significant differences (p ¡ 0.05), this is primarily an artifact of the massive sample size inherent to high-resolution, voxel-wise tomographic data. Crucially, the effect sizes for all morphological metrics were consistently trivial (*Cohen*^′^*sd <* 0.2). Therefore, while the influence of vasculature on local anisotropy and shape measures is statistically detectable, its actual magnitude and practical impact on the underlying white matter characterization are negligible.

**Fig. A5.**
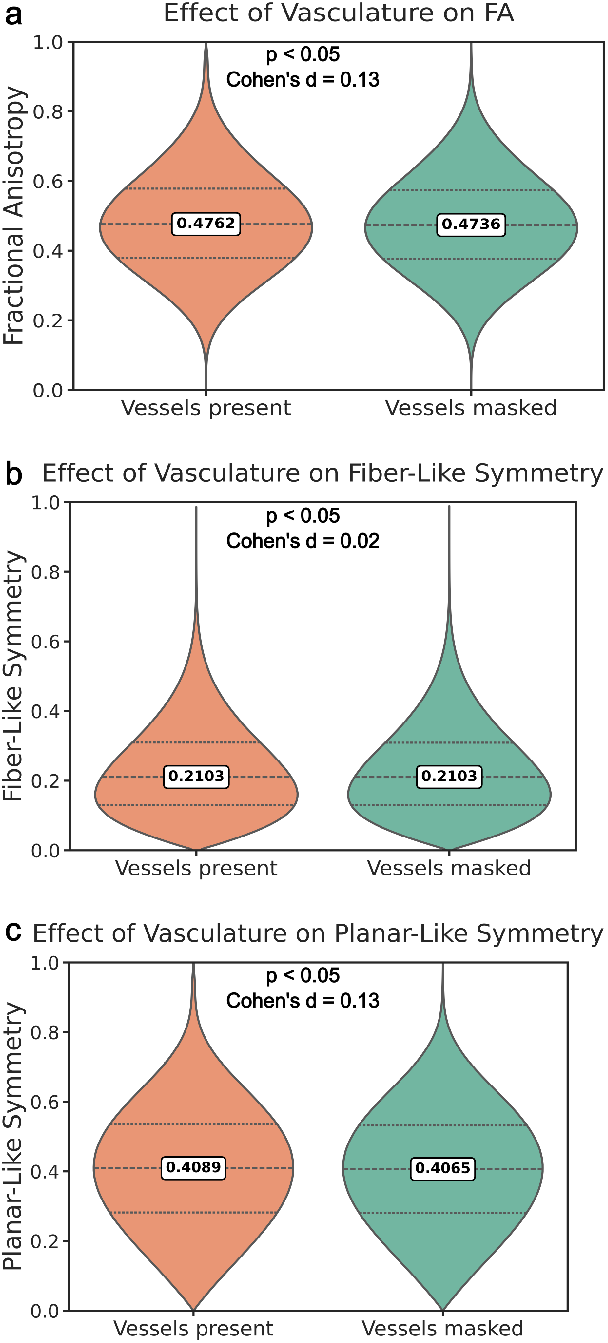
Impact of vascular masking on white matter anisotropy and shape metrics. Violin plots display the voxel-wise distributions of **(a)** Fractional Anisotropy (FA), **(b)** linearity, and **(c)** planarity computed with and without gradient-level vascular masking. While massive voxel counts drive statistically significant differences (p ¡ 0.05), the highly overlapping distributions and trivial effect sizes (Cohen’s d ¡ 0.2) demonstrate that vasculature has a negligible magnitude of effect on the overall characterization of white matter morphology.

### A.5 Effects of artifacts on HiP-CT STA based tractography

The use of Fomblin, a perfluoropolyether fluid routinely employed during ex vivo MRI to prevent tissue dehydration and minimize magnetic susceptibility artifacts, introduces specific challenges for correlative HiP-CT imaging. Despite extensive sample preparartion and washing protocols following the dMRI scans, residual fomblin can remain stubbornly trapped within deep sulci and vascular cavities **Figure A6 a, b**. Because fomblin has a vastly different X-ray attenuation coefficient profile and electron density compared to the surrounding fixed brain tissue, these residual droplets create sharp, high-contrast structural boundaries. During tomographic reconstruction, these sharp density transitions generate severe streak artifacts that propagate linearly across the adjacent tissue **Figure A6 c, e**. Because structure tensor analysis relies fundamentally on local spatial intensity gradients to estimate fiber orientation, these streak artifacts act as a massive confounding factor and are misinterpreted as highly coherent biological tissue. Consequently, this leads to the generation of dense, spurious streamlines that align with the artificial gradient trajectories **Figure A6 d, f**. These observations reinforce the necessity of careful HiP-CT sample preparation, reconstruction and quality control.

**Fig. A6.**
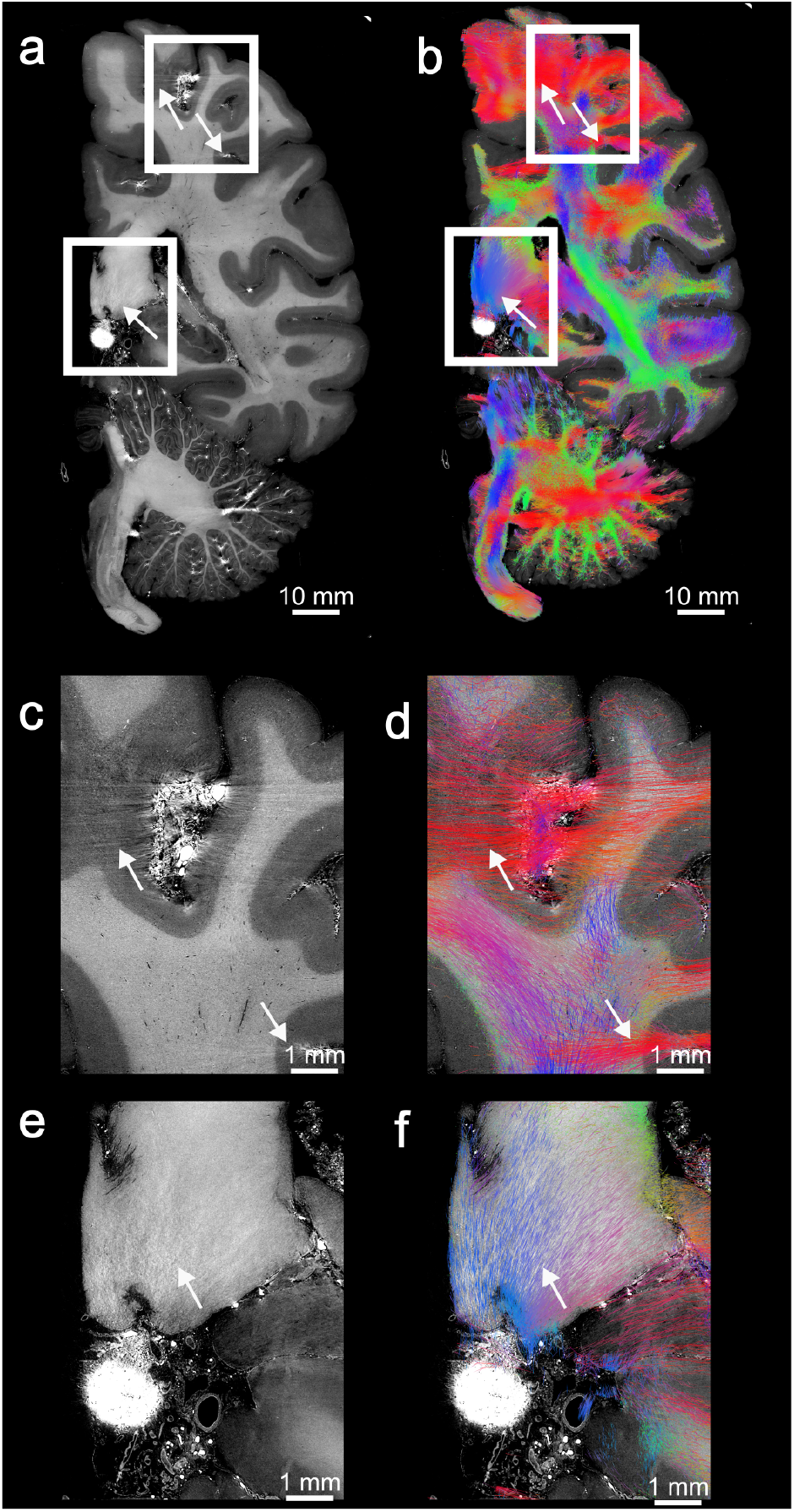
Impact of residual Fomblin and tomographic streak artifacts on HiP-CT structure tensor tractography. **(a, b)** Macroscopic coronal cross-sections of HiP-CT intensity (a) and corresponding structure tensor tractography (b). Highly attenuating Fomblin fluid, retained from prior ex vivo dMRI sample preparation, pools in macroscopic cavities and deep sulci. **(c, e)** Magnified regions revealing severe, high-frequency streak artifacts propagating from the dense fomblin deposits during tomographic reconstruction. **(d, f)** The corresponding structure tensor tractography for these magnified regions. Because the algorithm relies on local spatial derivatives, it mathematically misinterprets the strong, artificial intensity gradients of the streak artifacts as highly coherent biological tissue. This results in the generation of dense, spurious streamlines (white arrows) that align with the artifactual streaks, most notably the dense horizontal tracts in (d), locally obscuring and overwriting the true underlying white matter architecture.

### A.6 Beyond dMRI equivalent resolution

To demonstrate the scalable potential of the pipeline beyond matched dMRI resolution, we repeated the structure tensor analysis using 400 µm supervoxels instead of the 800 µm supervoxels used in the main text. The high resolution 15 µm field was first grouped into 400 µm supervoxels, after which spherical histograms were constructed and fitted with 8th-order spherical harmonics to obtain fODFs, exactly as described for the 800 µm case. All other tractography parameters remained identical. The resulting 400 µm tractogram are shown in Supplementary Figure A7. Reducing the supervoxel size preserved the overall bundle organization and global topological agreement with the 800 µm results while visually yielding denser, continuous and spatially confined streamlines.

**Fig. A7.**
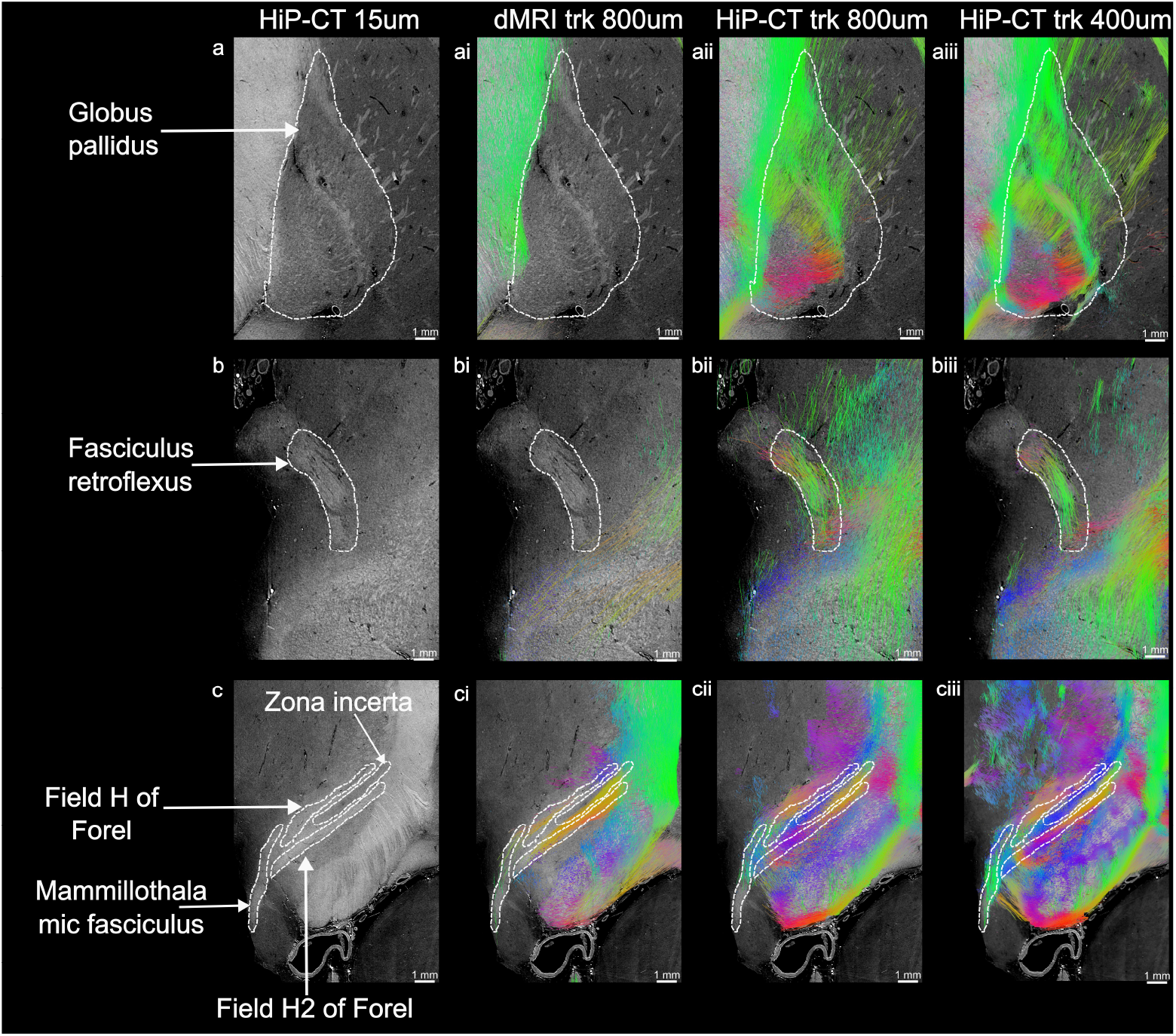
Comparison of the same deep-brain regions shown in Figure 4, now including HiP-CT-STA tractography performed with 400 µm supervoxels (rightmost column). Reducing the aggregation scale preserved the global topological organization of the major white matter bundles while minimizing partial volume effects yielding yielding denser, continuous and spatially confined streamlines. Streamlines are colored by local orientation (Red: left–right, Green: anterior–posterior, Blue: superior–inferior).

